# Discovery and 3D imaging of a novel ΔNp63-expressing basal cell type in human pancreatic ducts with implications in disease

**DOI:** 10.1101/2020.08.20.259317

**Authors:** Sandrina Martens, Mathias Van Bulck, Katarina Coolens, Hediel Madhloum, Farzad Esni, Gunter Leuckx, Harry Heimberg, Luc Bouwens, Patrick Jacquemin, Peter In’t Veld, Pierre Lefesvre, Francisco X. Real, Meritxell Rovira, Ilse Rooman

## Abstract

**Objective:** An aggressive basal-like molecular subtype of pancreatic ductal adenocarcinoma (PDAC) exists, driven by ΔNp63. In other epithelia, ΔNp63^+^ basal cells have stem cell capacity and can be at the origin of tumors. In the pancreas, basal cells have not been identified.

**Design:** We assessed basal cell markers in human and mouse pancreas, chronic pancreatitis and PDAC, and developed a 3D imaging protocol (FLIP-IT) to study sizeable samples at single cell resolution. We generated organoid cultures of ducts from Sox9-eGFP reporter mice.

**Results:** In normal human pancreas, rare ΔNp63^+^ cells exist in ducts that expand in chronic pancreatitis. ΔNp63^+^ cells express KRT19 and canonical basal markers (KRT5, KRT14 and S100A2) but lack markers of duct cells such as CA19.9 and SOX9. In addition, ΔNp63^+^ cells pertain to a niche of cells expressing gastrointestinal stem cell markers. 3D views of the ductal tree in formalin fixed paraffin embedded samples show that basal cells are localized on the basal membrane of medium to large ducts and expand as multilayer dome-like structures in chronic pancreatitis. In mice, ΔNp63 expression is induced when culturing organoids from Sox9-low ductal cells but could not be found in normal pancreas nor in models of pancreatitis or pancreatic cancer.

**Conclusion:** We discovered a novel ductal cell population in normal human pancreas similar to basal cells in other tissues. Using FLIP-IT, we provide unprecedented 3D visualization of these cells in archival clinical specimens. ΔNp63^+^ cells may play an important role in pancreatic tissue regeneration and cancer.

**SUMMARY BOX:** **What is already known about this subject?**

- ΔNp63 has a central role in determining the basal-like subtype of pancreatic ductal adenocarcinoma (PDAC).
- Different to other tissues with basal cancers, the normal pancreas reportedly does not contain (ΔNp63-expressing) basal cells.
- Current protocols face severe limitations for marker-based identification and 3D imaging of individual (rare) cells in archival pancreatic samples.

**What are the new findings?**

- We report a rare and atypical pancreatic duct cell that expresses ΔNp63, other basal cell markers and g.i. stem cell markers.
- The number of these basal cells increases in diseases such as chronic pancreatitis and pancreatic cancer.
- We provide an easy to implement protocol for 3D clearing and high-resolution imaging of sizeable samples of (fresh or FFPE) human pancreas or an entire mouse pancreas.
- Except after culturing medium to large ducts as organoids, we fail to detect basal cells in mouse experimental pancreatic models.

**How might it impact on clinical practice in the foreseeable future?**

- Extrapolating from knowledge in other organs, basal cells in the pancreas may have a stem cell/progenitor role, including in diseases such as (basal) pancreatic cancer.
- Use of the 3D imaging protocol in archival clinical specimens will allow unprecedented insights in pancreatic histopathology.
- For above mentioned diseases, we caution for findings in experimental mouse models that may not (fully) recapitulate the etiopathogenesis.

## INTRODUCTION

Pancreatic ductal adenocarcinoma (PDAC) is a cancer of high unmet need with rising incidence. In several large PDAC cohorts, a basal-like/squamous molecular subtype has been identified, having the worst prognosis (1). ΔNp63, an isoform of tumor protein P63 (2), was found a driver of this basal-like subtype (3). In many epithelial organs such as the bronchus, prostate, salivary gland, skin, breast and placenta (4), ΔNp63 is expressed by a specific cell population in normal ducts, located on the basement membrane, hence termed basal cells, and distinguished by specific markers among which cytokeratin (KRT) 5 and 14 (5). Furthermore, ΔNp63 is a well-studied key player in the development of stratified epithelium and an inhibitor of cell differentiation, crucial for stem cell renewal (6,7). Accordingly, basal cells are often described as progenitor cells, which are important in development, adult tissue homeostasis and repair mechanisms (5,8,9). On the other hand, ΔNp63 is a driver of poor prognostic basal subtypes of various cancers in basal cell containing epithelia (10). Several papers have reported that ΔNp63^+^ basal cells can be the cells of origin of the tumors (11,12). Despite having a ΔNp63-driven tumor, in the normal pancreas it is accepted that there is no expression of any P63 isoform (3,4,13).

Studying the spatial organization of cells at single cell level in the pancreas requires tissue clearing techniques together with fluorescent labelling strategies and 3D imaging of the cells in sizeable experimental and clinical samples (14,15). However, such approach faces several limitations especially when it comes to clinical specimens that are often formalin fixed and paraffin embedded (FFPE). Applying the current tissue clearing methods, such as CUBIC, CLARITY and DISCO, that found their origin in brain research and were mainly used to study macroscopic changes (16), provide suboptimal results when imaging pancreatic tissue, particularly when using light sheet fluorescence microscopy (LSFM) for achieving single cell resolution.

Here, we report a novel rare cell population that expresses ΔNp63, amongst other basal cell markers. We developed a clearing and multiplex 3D imaging protocol, circumventing current technical limitations. Using archival clinical samples, we assessed the pancreatic basal cells’ topography in normal pancreas and chronic pancreatitis. Contrary to human samples, we failed to detect this basal cell population in the normal mouse pancreas and in a wide variety of commonly used experimental *in vivo* models for pancreatic cancer. Yet, a ΔNp63-expressing cell population was identified in organoid cultures from medium to large size ducts indicating that this phenotype can be acquired by mouse ductal cells.

The discovery of a basal cell population in the human pancreas raises important conceptual questions about their developmental origin, fate, and role in regeneration and disease, including the basal-type PDAC with which it shares many markers.

## RESULTS

### Normal human pancreatic ducts harbor a rare ΔNp63^+^ cell population

We confirmed previous reports (1,3) that a subset of PDAC (46 out of 141) were positive for ΔNp63 (S Fig 1). Some of these tumors displayed sparse positivity for ΔNp63, while other tumors were almost completely positive and had the strongest staining intensity. The latter included adenosquamous and less differentiated tumors (S Fig 1). ΔNp63^+^ cells were located at the basal side of well-differentiated tumoral ducts, while in less differentiated tumors, ΔNp63^+^ cells were randomly localized.

Interestingly, in 6 out of 10 samples with presumed normal tissue adjacent to the tumor, ΔNp63^+^ cells could also be found in the lining of normal ducts.

Hence, we assessed pancreatic tissue of healthy organ donors who had no history of pancreatic disease. When one random section (67.20 ± 6.8 mm^2^) was assessed, we detected strong but rare ΔNp63 expression in ducts (Fig 1A) in 53 of 113 donors (45%). This was confirmed by RNA *in situ* hybridization for ΔNp63 and by immunofluorescence (IF) with an anti-P63 antibody that detects all isoforms (S Fig 2). In the normal tissue, regardless of whether it was from head or tail region, ΔNp63^+^ cells were detected as single cells in the basal lining of a duct, in small clusters around ducts, as a combination of single cells and a cluster, or very rarely as apparently single cells among acini (Fig 1B-E), although these observations are limited by interpreting 2D sections.

**Figure 1:**
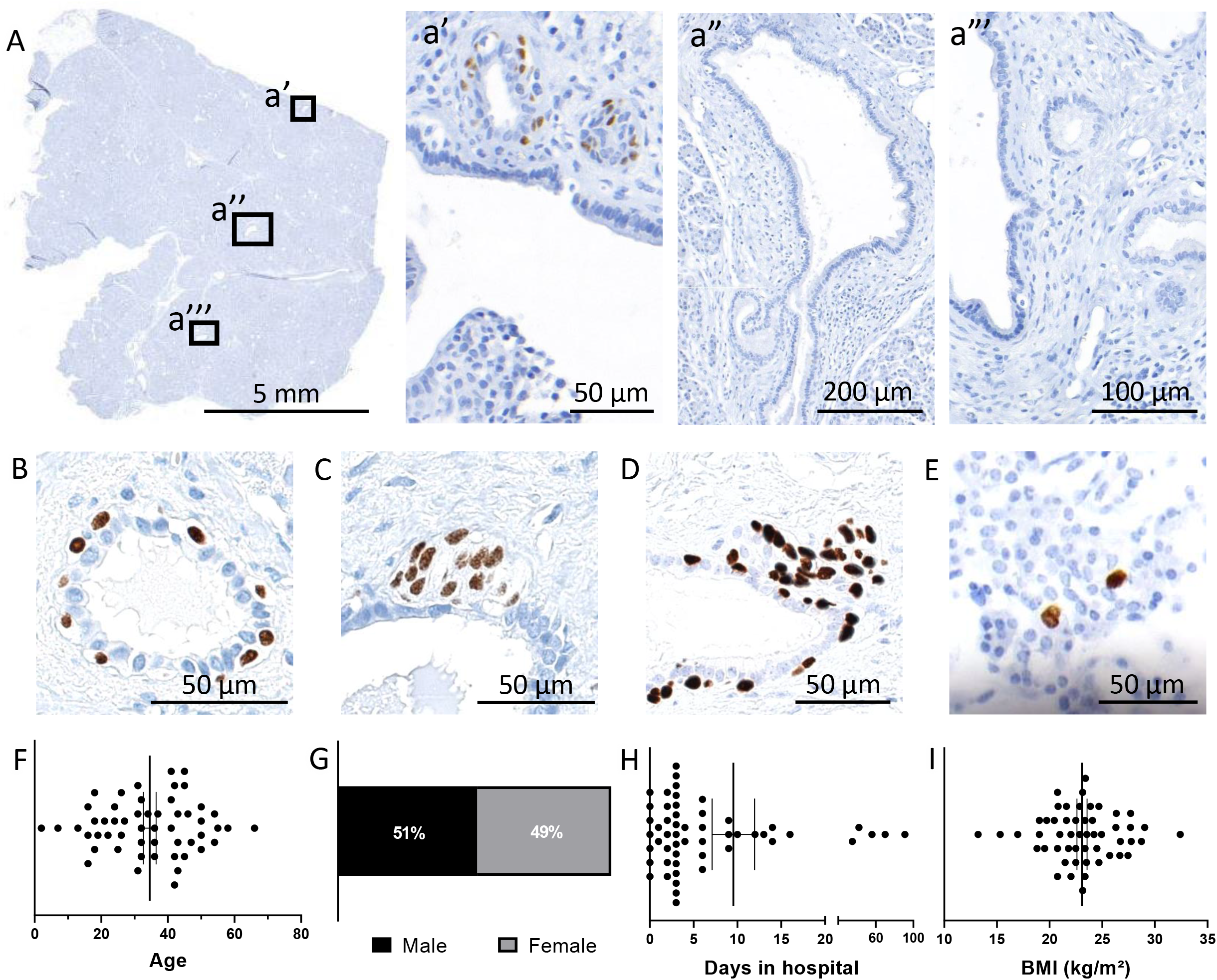
ΔNp63^+^ cells are a rare ductal cell population in the normal human pancreas. (A) ΔNp63 staining in one representative section of healthy donor pancreata (n=114): Three ductal areas are shown, of which one (a’) contains some positive cells, and two other areas (a” and a’’’) are negative; (B-E) Location of ΔNp63^+^ cells throughout a healthy pancreas: Cells can be located basally within a duct (B), they can form a small cluster near a duct (C) or there can be both groups and single cells combined (D). ΔNp63^+^ cells can rarely be found within acinar tissue (E). (F-I) Characteristics of all human pancreas donors with ΔNp63 detected in a section (n=53): (F) Age, (G) Gender, (H) Days spent in the intensive care unit, (I) BMI. Mean ± SEM is shown.

Of the ΔNp63-positive samples, we evaluated several donor characteristics (S Table 1) and found a random distribution in age (Fig 1F), gender (Fig 1G), the time in the intensive care unit (Fig 1H), body mass index (BMI) (Fig 1I) and fixation protocol of the tissue (not shown). We also compared donor characteristics between ΔNp63-positive and negative samples and found no difference (S Fig 3), indicating that the presence of ΔNp63^+^ cells did not associate with specific donor characteristics. Indeed, when analyzing more than one FFPE block (here n=4 donors with two to ten FFPE blocks available), at least one block harbored ΔNp63^+^ cells. This suggested that ΔNp63 cells could likely be found in the pancreas of any donor if sufficient material was analyzed.

We thus report for the first time ΔNp63-expressing cells in the adult pancreas of subjects without pancreatic disease.

### ΔNp63^+^ cells expand in chronic pancreatitis

Next, we assessed samples of patients with chronic pancreatitis, a condition with known expansion of ducts and an established risk factor for the development of PDAC (17). When analyzing one section per patient (228.3 ± 17.0 mm^2^), 9 out of 11 (82%) chronic pancreatitis samples were positive for ΔNp63 (Fig 2A). ΔNp63^+^ cells were usually grouped in large numbers near ducts and were especially located in the ductal lining of cysts (Fig 2A a’).

**Figure 2:**
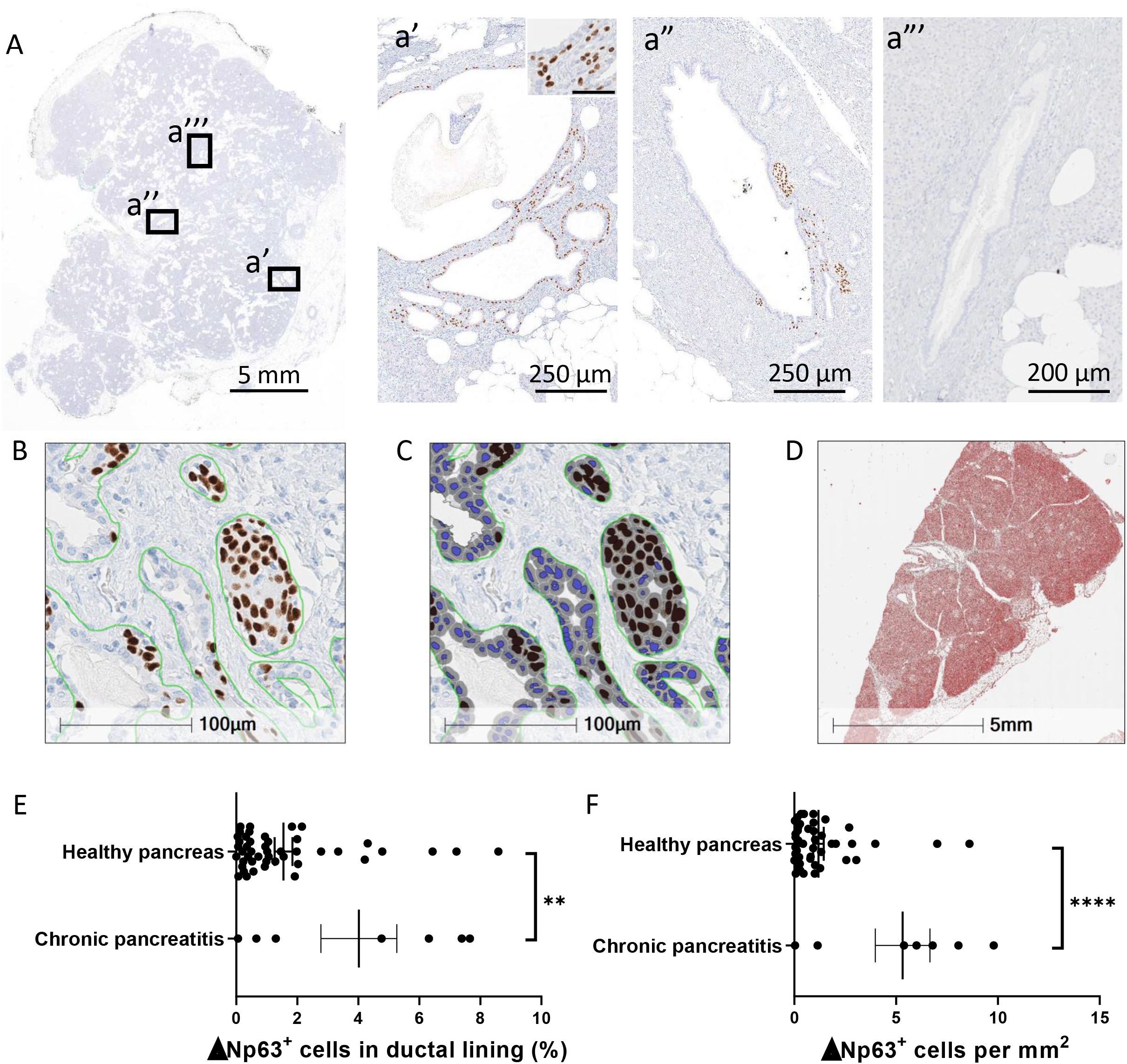
ΔNp63^+^ cells expand significantly in chronic pancreatitis. (A) ΔNp63 staining in one representative section of human chronic pancreatitis (n=11). Three ductal areas are shown, of which two (a’ and a’’) contain a high number of positive cells, and one area (a’’’) is negative. An inset is shown in figure a’. (B-D) Quantification of ΔNp63 cells done with HALO; (B) shows annotation of all ducts, (C) shows quantification of this annotation and (D) shows area quantification, tissue is indicated in red. (E) Percentage of cells within the ductal lining in normal pancreas (n=46) and chronic pancreatitis (n=7) (**P= 0,044); (F) Quantification of number of ΔNp63 cells per mm^2^ in normal pancreas (n=46) and chronic pancreatitis (n=7) (****P<0,0001). Mean ± SEM is shown.

In pancreas from healthy patients and patients with chronic pancreatitis, we annotated all ducts (Fig 2B) and analyzed the number of duct cells and ΔNP63-positive cells (Fig 2C) and the size of each section (Fig 2D). In healthy pancreas, 1.6% of all cells within the ductal lining were ΔNp63^+^ (Fig 2E) averaging 1.2 cells per mm^2^ of pancreatic tissue (Fig 2F). The occurrence of ΔNp63^+^ cells in chronic pancreatitis was significantly higher (4.0%, 5.3 cells per mm^2^), albeit with large variability among the patients (Fig 2E, F).

In conclusion, we find a significant expansion in the number of ΔNp63-expressing cells in chronic pancreatitis patients.

### ΔNp63^+^ cells are distinct from normal pancreatic duct cells and have typical basal cell markers

To determine the identity of ΔNp63-expressing cells, we analyzed the expression of an epithelial cell marker (E-Cadherin), canonical pancreatic ductal cell markers (KRT19, CA19.9, SOX9, HNF1β and KRT7), markers of pancreatic duct glands (PDG) (MUC6), a suggested proliferative (and Ki67-positive) stem cell niche (18), as well as basal cell markers reported in other tissues, i.e. expression of KRT5, KRT14 and S100A2 (5,19,20) and basal positioning, pale cytoplasm and nuclei (21). In the breast, myoepithelial cells have been shown to express ΔNp63 as well as myogenic markers (22). So, we also analyzed expression of calponin and alpha smooth muscle actin (aSMA).

ΔNp63^+^ cells were positive for E-cadherin (S fig 4) and KRT19 (Fig 3A) but not for CA19.9. Nuclear expression of SOX9 and HNF1β was absent, while the neighboring ΔNp63^−^ duct cells stained positive (Fig 2C, D). ΔNp63^+^ cells also lacked ductal KRT7 (Fig 2E). Finally, ΔNp63^+^ cells were MUC6 and Ki67 negative, while nearby cells of PDG were clearly positive (S fig 5).

**Figure 3:**
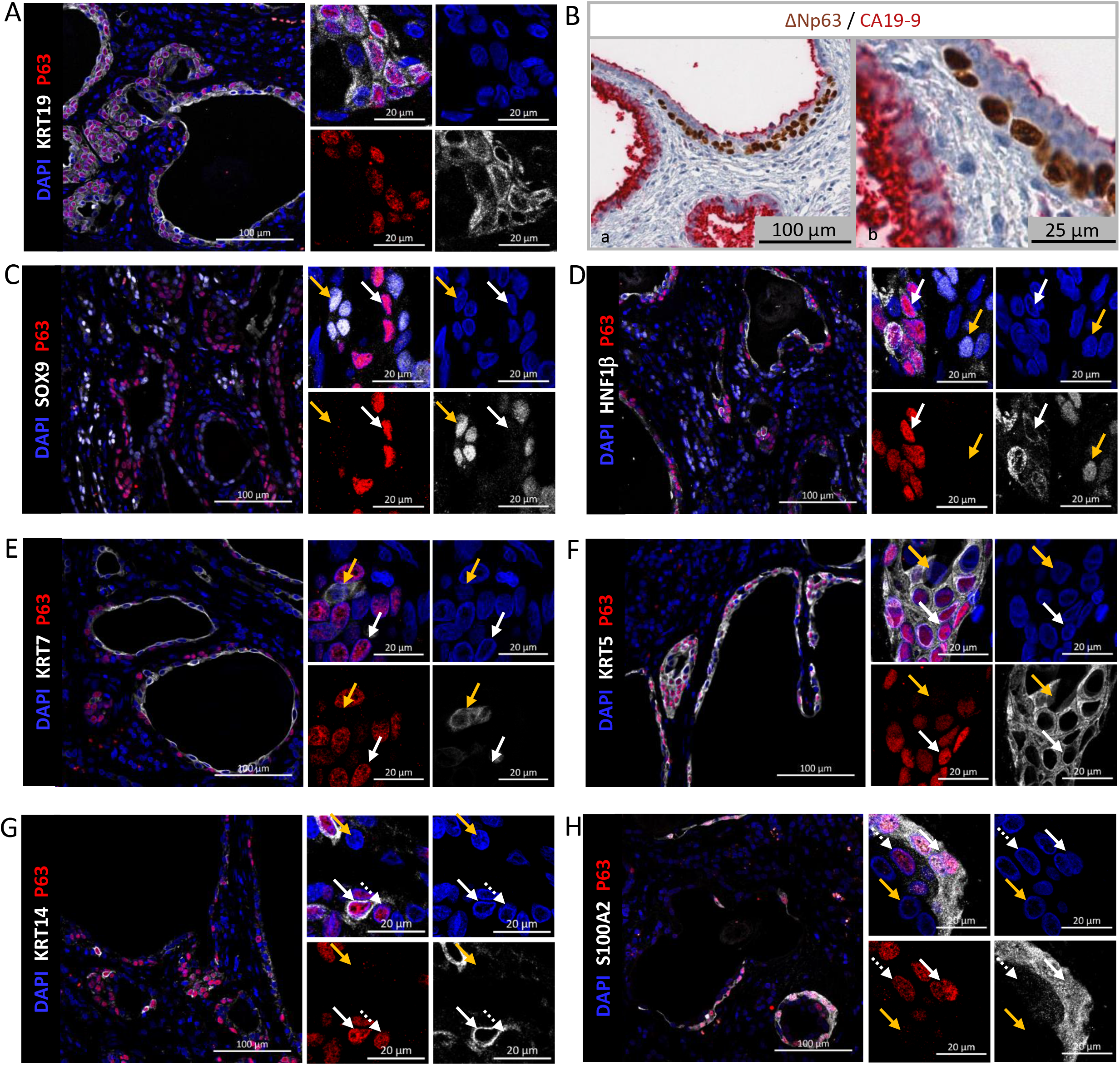
ΔNp63^+^ cells are epithelial cells expressing canonical basal markers and lacking typical ductal markers. (A) IF for KRT19 (white) and P63 (red), showing co-expression; (B) IHC for ΔNp63 (brown) and CA19.9 (red). CA19.9 expression is absent in ΔNp63^+^ cells and weaker in the duct lumen (see inset); (C) IF for SOX9 (white) and P63 (red). SOX9^+^ cells are indicated with an orange arrow, while ΔNp63^+^ cells are indicated with a white arrow; (D) IF for HNF1β (white) and P63 (red). HNF1β^+^ cells are indicated with an orange arrow, while P63^+^ cells are indicated with a white arrow; (E) IF for KRT7 (white) and P63 (red). White arrow indicates a ΔNp63^+^ cell, while the orange arrow indicates a KRT7^+^ cell; (F) IF for P63 (red) and KRT5 (white). White arrow indicates a P63^+^ cell, orange arrow indicates a P63^−^ cell; (G) IF for P63 (red) and KRT14 (white). Full white arrow indicates a ΔNp63^+^KRT14^+^ cell, dotted white arrow indicates a ΔNp63^+^KRT14^−^ cell. Orange arrow indicates ΔNp63^−^ cell; (H) IF for S100A2 (white) and P63 (red). Full white arrow indicates a ΔNp63^+^S100A2^+^ cell, dotted white arrow indicates a ΔNp63^+^S100A2^−^ cell. Orange arrow indicates ΔNp63^−^ cell.

On a HES staining (S fig 6), ΔNp63^+^ cells often presented with a basal location, a paler cytoplasm and paler nuclei, compared to ΔNp63^−^ cells in the duct. All ΔNp63^+^ cells strongly expressed the basal cell marker KRT5 (Fig 3F) with a subset also expressing KRT14 (Fig 3G). A large subset of the ΔNp63^+^ cells stained positive for S100A2, a direct transcriptional target of ΔNp63 (23) (Fig 3H), as do basal cells of prostate and airway epithelium (19,20). The pancreatic ΔNp63^+^ cells did not express aSMA or calponin (S Fig 7).

In conclusion, pancreatic ΔNp63-expressing cells have a phenotype like that of canonical basal cells from other tissues and represent a distinct population of normal ductal cells.

### Ducts containing ΔNp63^+^ cells express gastrointestinal stem cell markers

In other epithelia, basal cells are thought to be progenitors of more differentiated epithelial cells. Therefore, we investigated whether the pancreatic ΔNp63^+^ cells also harbored characteristics of gastrointestinal (GI) stem cells such as DCLK1, CD142 and OLFM4 (24) (25), (26) or, more general, of pluripotent stem cells (NANOG, OCT4).

Most basal ΔNp63^+^ cells stained positive with anti-DCLK1 (Fig 4A). Singular DCLK1^+^ P63^−^ cells were also observed, confirming previous reports in pancreas (27,28) (Fig 4B). ΔNp63^+^ basal cells and some neighboring luminal cells, specifically of ducts containing ΔNp63^+^ cells, expressed CD142 while CD142 was not found in ducts lacking ΔNp63^+^cells (Fig 4C,D). OLFM4 was absent in ΔNp63^+^ cells themselves and rarely expressed in cells neighboring ΔNp63^+^ cells (Fig 4E). However, we found it exclusively in the lumen of ducts containing ΔNp63^+^ cells (Fig 4E, asterisk and F). OLFM4 was reported in bile (29) but here the intraluminal product was negative for the Hall’s bilirubin staining (not shown), hence we discarded it as being bile. ΔNp63^+^ cells did not express NANOG or OCT4, contrary to positive control samples (S fig 8).

**Figure 4:**
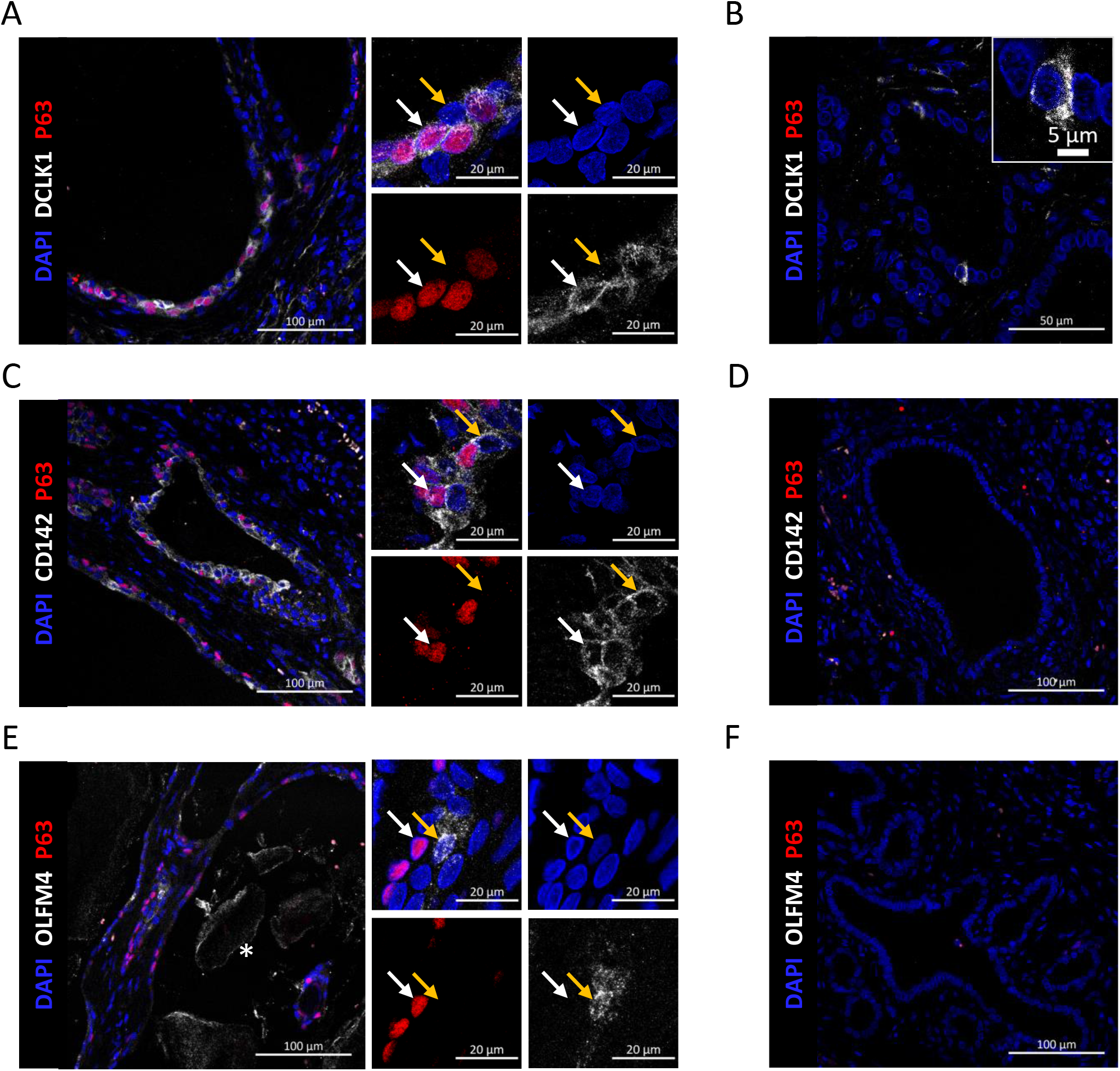
Gastrointestinal stem cell markers are expressed in ΔNp63^+^ ductal structures. (A) IF for DCLK1 (white) and P63 (red). P63^+^ cells express DCLK1, but are not like tuft cells (B) that are solitary cells with apical microvilli and no expression of P63; (C) IF for P63 (red) and CD142 (white). P63^+^ cells and the cells located on the apical side of P63^+^ cells express CD142, unlike ducts without P63^+^ cells (D); (E) IF for P63 (red) and OLFM4 (white). Ducts that contain P63^+^ cells secrete OLFM4 into their lumen (indicated with asterisk) and rarely contain OLFM4^+^ cells, unlike ducts that do not contain P63^+^ cells (F).

In summary, ducts containing pancreatic basal cells show expression of GI stem cell markers, either in the basal cells, in the juxtaposed cells or both, suggesting the basal cells pertaining to a stem cell niche.

### FLIP-IT allowed 3D visualization of pancreatic basal cells in archival FFPE tissue

To visualize the ΔNp63^+^ cell population in a human pancreas at single cell resolution in 3D, we require high magnification and high numerical aperture objectives in conjunction with highly cleared samples and preservation of fluorescence intensity. So far, imaging on a high magnification (≥ 20x) can only be achieved with lengthy confocal or two-photon microscopy thus limiting the scanning capabilities, increasing the scanning time and experiencing loss of signal due to photobleaching (30,31).

Hence, we developed a protocol for <u>F</u>luorescence <u>L</u>ight sheet microscopic <u>I</u>maging of <u>P</u>araffin-embedded or <u>I</u>ntact <u>T</u>issue (FLIP-IT) for which we optimized the permeabilization and delipidation processes. For the RI matching, we used CUBIC-R and ethyl cinnamate (ECi). FLIP-IT enabled us to assess FFPE and intact (fresh) tissues from patients and mice (Fig 5A and S Fig 9). The process from clearing to imaging of FFPE punches could be completed in less than two weeks, which is a significant reduction in processing time compared to published methods (16). Bulk processing of samples was possible as ECi preserved the fluorescence signal for weeks to months.

**Figure 5:**
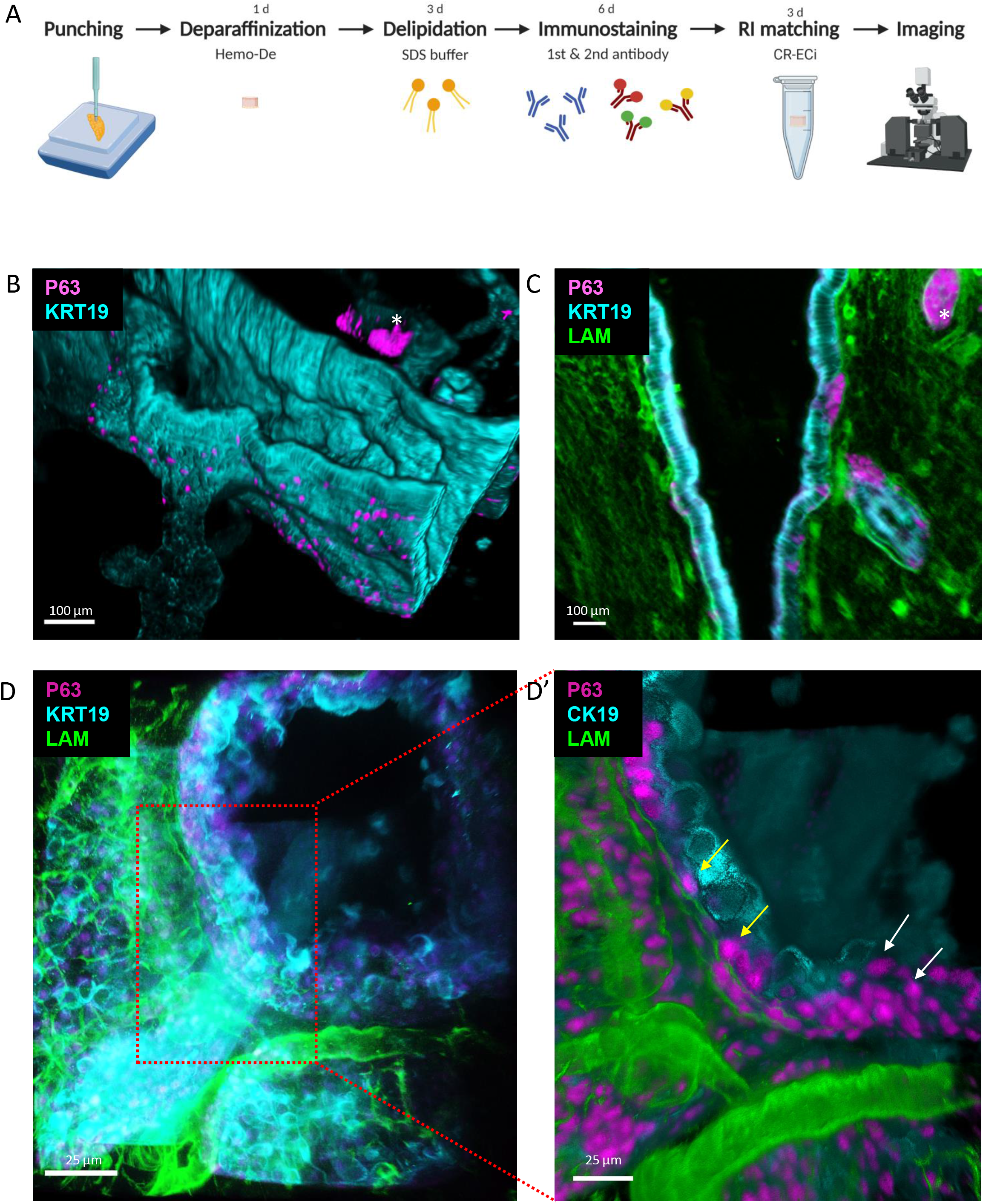
FLIP-IT and high resolution (HR)-Light sheet Fluorescence Microscopic (LSFM) reveal the 3D location of P63^+^ cells in normal human pancreas and chronic pancreatitis FFPE sample. (A) FLIP-IT protocol overview processing of human archival FFPE samples; (B) Overview 3D volume rendering of a large duct system (cyan) with P63^+^ cells (pink) in normal human pancreas. Objective 20x, zoom 0.36. Scale bar corresponds to 100μm; (C) Z-plane clipping of B with KRT19 (cyan), P63 (pink) and Laminin (green). Asterisk indicates reference structure in B. Objective 20x, zoom 0.36. Scale bar corresponds to 100μm; (D) HR-LSFM 3D rendering of a dome positive for P63 (pink) in chronic pancreatitis. Objective 20x, zoom 1. Scale bar corresponds to 25μm. (D’) Inset from D, HR-LSFM 3D volume rendering of D in chronic pancreatitis. Yellow arrows indicate P63^+^ (pink) cells in contact with the basal membrane (green). White arrows indicate p63^+^ not in contact with basal membrane. Objective 20x, zoom 1. Scale bar corresponds to 25μm. n≥2

Rare P63^+^ cells could be found as solitary cells in or as clusters attached to ducts with a minimal ductal diameter of 20 micron (Fig 5B), confirming the analyses of 2D sections. Co-staining with laminin showed that solitary P63^+^ cells lie between the basal lamina and the luminal cells of large ducts. The P63^+^ clusters also associated to a basal lamina (Fig 5C). In chronic pancreatitis, we observed KRT19^+^ domes of multiple cell layers around a lumen, where only the basally located P63^+^ cells were touching the basal membrane (Fig 5D,D’).

3D image rendering allowed for straightforward identification of KRT5^+^ KRT7^−^ basal cells within the pancreatic tissue (video 1, Fig 6A). Small clusters of round basal cells in normal pancreatic tissue (n=4 normal human pancreas, one punch analyzed in each) grouped around a small lumen (Fig 6C,C’ and S fig 7). In chronic pancreatitis patients (n=2 chronic pancreatitis patients, two punches for each analyzed) (video 1, Fig 6B), we found large dome-like clusters associated to cystic ducts that were already apparent from the overview 3D rendering, with KRT5^+^ cells located on the outside of the domes and KRT7^+^ cells lining the luminal side (Fig 6D,D’). 3D measurements of sphericity and volume at single cell level, that could not be measured in 2D sections, demonstrated that the domes consisted of flatter cells and that the cell volume of the KRT5+ cells had also increased in chronic pancreatitis compared to their equivalent in the normal human pancreas (S Fig 10).

**Figure 6:**
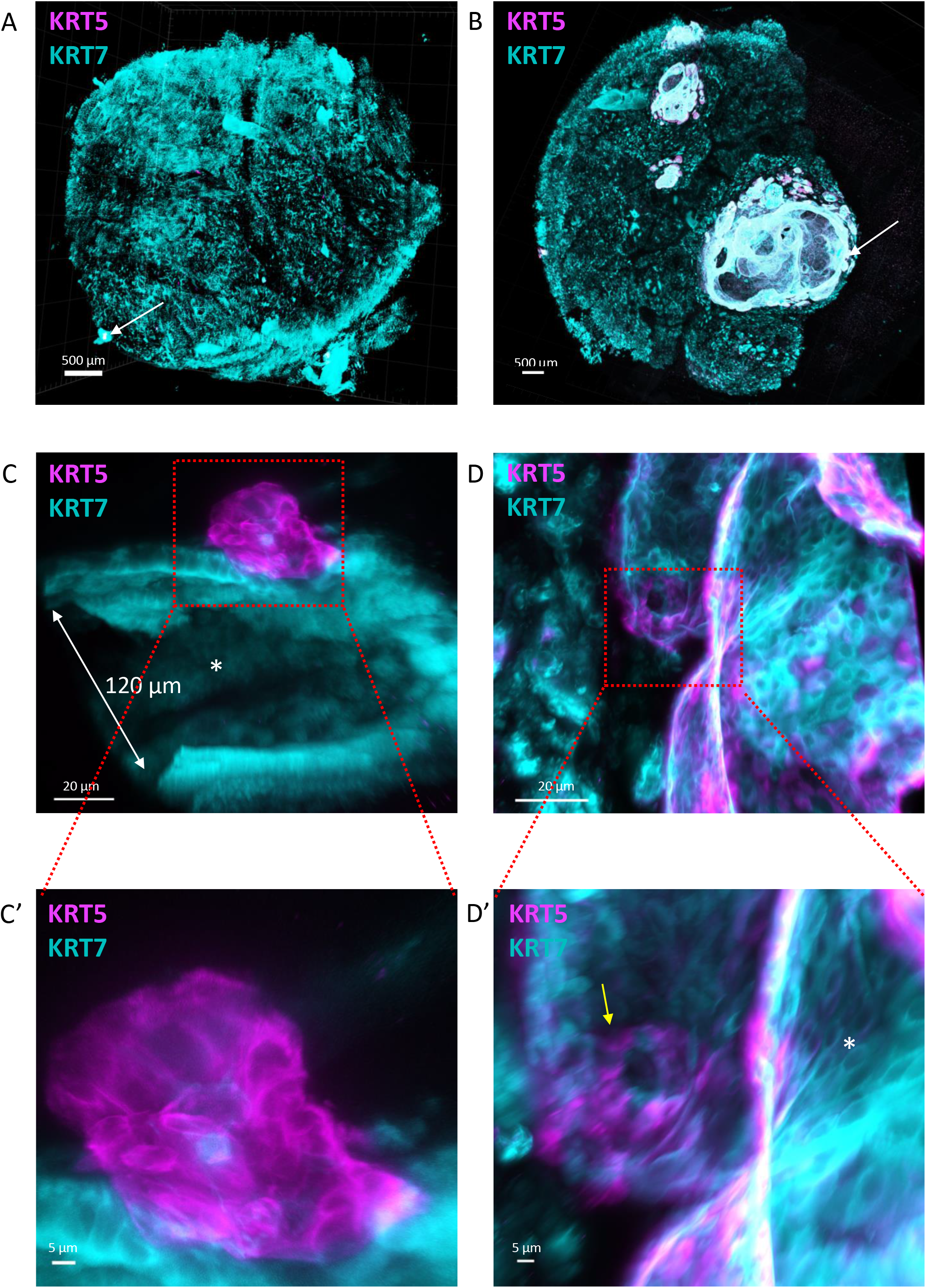
3D imaging of KRT5-positive basal cells in normal pancreas and expansion as dome-like structures in chronic pancreatitis. (A) Overview 3D rendering of normal human pancreas and (B) chronic pancreatitis from FFPE blocks stained for KRT5 (pink) and KRT7 (cyan). Arrows indicate magnified regions in (C) and (D) respectively for normal pancreas and chronic pancreatitis. Objective 20x, zoom 0.36. Scale bar corresponds to 500μm; (C) HR-LSFM of a KRT5^+^ (pink) dome on a large KRT7^+^ (cyan) duct with lumen diameter 120μm at widest point in normal pancreas. Asterisk indicates lumen. Objective 20x, zoom 2.5. Scale bar corresponds to 20μm; (D) HR-LSFM of dome wall showing its constitution in chronic pancreatitis. Objective 20x, zoom 2.5. Scale bar corresponds to 20μm; (C’) inset from C showing in great detail cellular structure of KRT5^+^ (pink) cells in normal pancreas. Objective 20x, zoom 2.5. Scale bar corresponds to 5μm; (D’) HR-LSFM of dome wall showing flat KRT5^+^ (pink) and KRT7^+^ (cyan) cells intercalated (yellow arrow) and KRT5^+^ (pink) cells lining the exterior of the dome wall. Asterisk indicates lumen. Objective 20x, zoom 2.5. Scale bar corresponds to 5μm. n≥2

Thus, leveraging on a novel imaging protocol, FLIP-IT, that is widely applicable, we established the spatial distribution and morphometric features of individual pancreatic basal cells within the ductal tree, in archival samples of normal human pancreas and chronic pancreatitis.

### ΔNp63 expression is undetectable in commonly used experimental mouse models of pancreatic disease but is induced in organoid cultures

We re-assessed ΔNp63 expression in mouse pancreas, since in single-cell RNA sequencing (scRNAseq) studies ΔNp63 in the pancreas was not reported (S Table 2). In contrast to the mouse mammary gland and skin (S Fig 11 A,B), ΔNp63^+^ cells were undetectable in the healthy pancreas, also when specifically examining the larger ducts such as the main pancreatic duct and the common bile duct (Fig 7A). Similarly, we did not find ΔNp63^+^ cells in pancreatic samples from pregnant, postpartum, neonatal and aged mice. To extend these findings to pathological conditions, we analyzed experimental models of chronic pancreatitis (caerulein-treated or pancreatic duct-ligated Kras WT mice and caerulein-treated Kras^G12D^ (KC) mice (32)). ΔNp63 expression was undetectable here as well as in tumor samples of Kras and p53 mutant KPC mice that recapitulate the development of PDAC (32). All these samples were also negative for KRT5 and KRT14 (not shown). 3D imaging of a whole mouse pancreas, including the main pancreatic duct (S Fig 12) confirmed the absence of KRT5^+^ cells. In addition, a KRT14-eGFP reporter mouse line confirmed the absence of expression of KRT14 when assessing for GFP or staining for KRT14 in the pancreas (whereas basal cells from the skin were positive, not shown).

**Figure 7:**
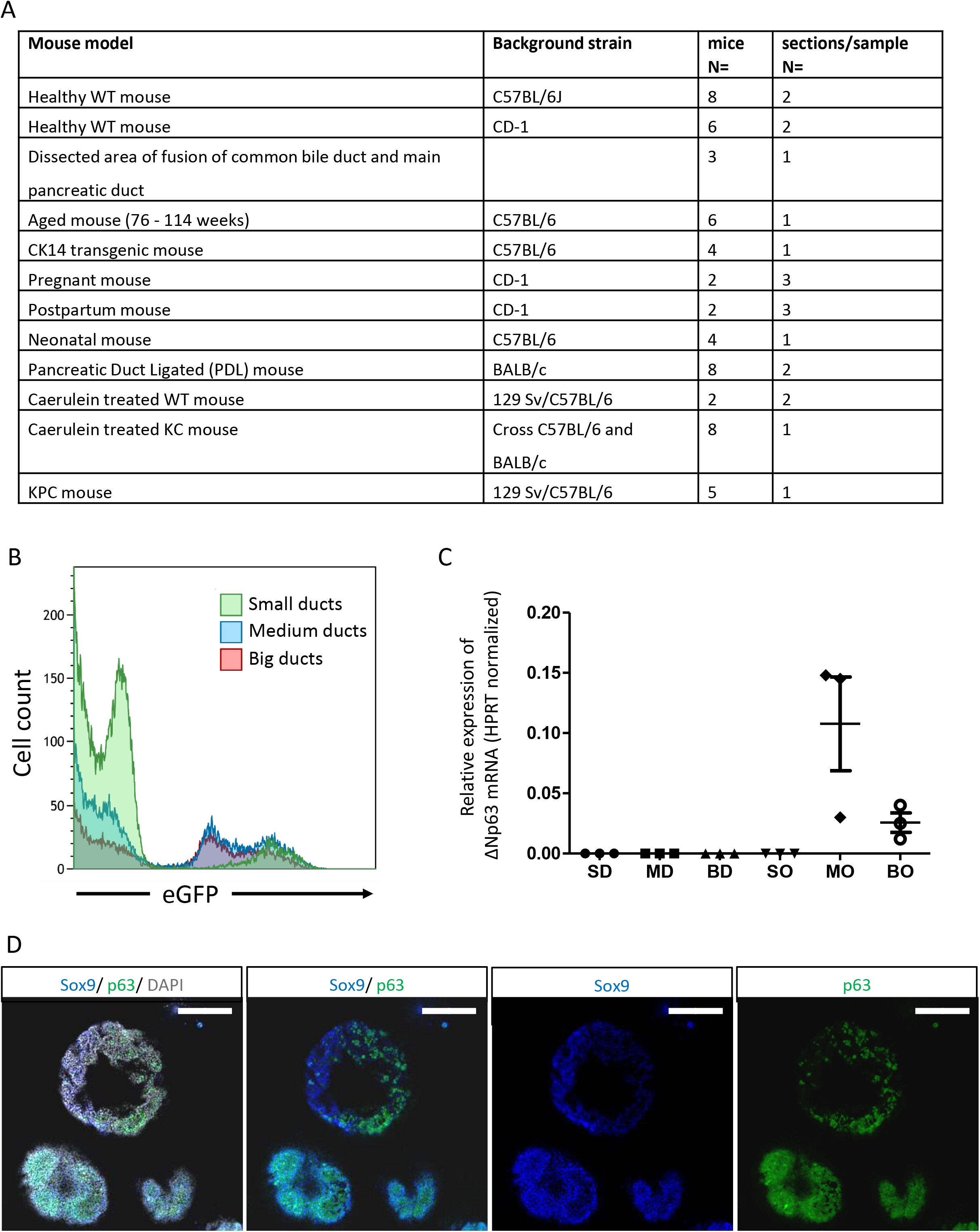
ΔNp63 can be induced in mouse pancreatic organoids but is undetectable in normal and experimental mouse disease models. (A) Table summarizing mouse models used to investigate ΔNp63 expression in rodent pancreas. IHC stainings for ΔNp63, KRT14 and KRT5. Background strain is indicated for each separate mouse model. Number of mice analyzed and sections analyzed per sample are indicated. (B) Representative flow cytometry plot of ductal cells from Sox9:GFP transgenic mouse line, showing different levels of Sox9:GFP expression that inversely correlate with the size of the ducts. (C) Bar plot indicates ΔNp63 mRNA levels normalized by HPRT of mouse ducts of different sizes (BD=big duct, MD=medium duct and SM= small duct) and organoids derived of the aforementioned ducts (organoids derived from big ducts = BO, medium ducts = MO and small ducts = SO). Error bars indicate SD of 3 independent experiments. (D) Representative immunofluorescence image of P63 (green), Sox9 (blue) and DAPI (gray) in organoids derived from big-medium size ducts. Scale bar = 200 μm.

We then investigated whether ΔNp63 expression is activated in murine pancreatic epithelial cells upon organoid formation, due to the induction of a progenitor cell-like phenotype in culture that could favor ΔNp63 expression. Thus, we used ductal cell digests from *Sox9*-eGFP reporter mice (33). These preparations showed intrinsic heterogeneous Sox9 expression, larger ducts having lower Sox9 expression (Fig 7B). Cells fractions were produced according to eGFP expression, corresponding to the size of the ducts. None of the fresh cell fractions showed detectable levels of ΔNp63 by real time qPCR (Fig 7C), confirming the tissue analyses (Fig 7A). Upon organoid culture, cells from Sox9-low medium to big size ducts showed an upregulation of ΔNp63 mRNA (Fig 7C). Whole mount stainings of organoids derived from low SOX9-expressing ductal cells showed a heterogeneous pattern of expression of ΔNp63 protein levels that correlated with the lowest levels of SOX9 (Fig 7D), reminiscent of the findings in normal human pancreas, where we failed to detect SOX9 in ΔNp63^+^ cells.

In conclusion, while normal mouse pancreas and models of pancreatitis and cancer do not show basal cells, organoid cultures from low SOX9-expressing pancreatic ducts can re-activate ΔNp63 expression, similar to that identified in human pancreata.

## DISCUSSION

Despite a strong consensus on the existence of a basal-like molecular subtype of PDAC driven by ΔNp63 expression (3), it is widely accepted that ΔNp63-expressing cells do not exist in healthy human and mouse pancreas (3,4,13). In this study, we provide compelling evidence of a ΔNp63^+^ cell population in the pancreas of individuals without a history of pancreatic disease. The lack of association with a large number of socio-demographic and clinical parameters strongly suggests that this cell fate is a constitutive feature of normal pancreatic differentiation.

The location of ΔNp63^+^ cells between the basal membrane and the luminal duct cells and their expression of KRT5, KRT14 and S100A2 support the idea that these cells represent the pancreatic counterpart of basal cells from other epithelial tissues such as the skin and lung. Like airway basal cells, pancreatic ΔNp63^+^ cells can be either KRT14^+^ or KRT14^−^, while they are all KRT5^+^ (34). Interestingly, KRT5 and KRT14 expression was previously reported in the pancreas (35) in less than 5% of ductal cells, which corresponds to our findings. Remarkably, studies using transmission electron microscopy reported a basally located cell type in interlobular ducts and the main pancreatic duct in human and rat pancreas, which were suggested to be a source of new ductal cells (36,37). Previous studies were however limited by the lack of access to large collections of pancreata from subjects without pancreatic disease and the inappropriate tools used to identify basal cells. Altogether, our findings call for a re-evaluation of the concept that the pancreatic duct is a homogenous “simple epithelium”, as established in classical histology textbooks.

The ΔNp63^+^ basal cells of the pancreas are situated in the ductal tree and express KRT19(38)(36)(38)(38) but the absence of CA19.9, SOX9 and HNF1β indicate that these cells represent a novel pancreatic ductal cell type. scRNAseq has been used in recent years to identify novel cell types in tissues. However, this strategy has so far failed to provide evidence of a basal cell type in the normal pancreas. A recent paper on human duct cell heterogeneity (39) did not report a group of cells with basal cell characteristics, possibly because only ALK3-positive cells were analyzed and ALK3 itself has not been reported in basal cells (19). Other single cell sequencing papers of human pancreatic tissue (40,41) did not pick up a (ΔN)p63 expressing cell population either. The rarity of ΔNp63^+^ cells, their restricted distribution along the ductal tree, and the shallowness of current scRNAseq technologies likely account for the failure of this powerful technology to identify the novel basal cell type that we report here. In one single cell sequencing dataset, we detected rare cells expressing ΔNp63 in samples of patients with type 2 diabetes (42). This may point to their increased presence in disease, concurring with our observations in chronic pancreatitis samples and in cancer.

To study the features of ΔNp63^+^ cells in normal pancreas and the changes associated with disease, we established a 3D imaging pipeline that allowed for the first time assessing cubic millimeters of a clinical sample. In the case of mouse pancreas, the whole organ could be imaged. Such 3D imaging protocol could have *a priori* revealed the existence of this cell type but was likely never undertaken because of several limitations (S Fig 9). Using FLIP-IT with punches of archival clinical samples, we visualized the ductal tree at single cell resolution and confirmed the existence of rare P63^+^/KRT5^+^ cells at the basal membrane in KRT19^+^/KRT7^+^ ducts with a minimal diameter of 20 μm. In chronic pancreatitis, the basal cells were organized as larger multilayer dome structures where they stretched out intercalated in between ductal cells in the outer layer and touching the basal membrane. The domes could reach sizes of cubic millimeter order. Thus FLIP-IT allows for unprecedented 3D static views and videos of (rare) pancreatic cells stained for markers of choice. FLIP-IT will be widely applicable for studies in which spatial insights are important and will endow researchers with a wealth of information in terms of (pancreatic) histopathology, in contrast to 2D assessments that only provide partial insights, limited in size and depth, and can lead to misconceptions because of sectioning artifacts. We envision that the 3D approach at single cell resolution can inform about the exact positioning of, for example, stromal cell types versus tumor epithelium. Another example of added value of FLIP-IT would come when studying the co-existence of different molecular subtypes within a tumor. We envision that the technical developments we report on will also allow expanding current technology to 3D-spatial transcriptomics.

In contrast to the solid evidence in human tissue, we failed to detect any basal cell marker in the adult mouse pancreas, including a series of *in vivo* experimental disease models, which in other organs activate the basal cell population (43,44). Bearing in mind that SOX9 levels in basal cells of human pancreatic ducts were undetectable, we assessed mouse pancreatic duct cells according to their variable Sox9 expression. Indeed, culturing the Sox9-low expressing medium and larger size ducts in stem cell-favoring organoid conditions induced ΔNp63 expression. This illustrated an inherent potential of mouse pancreatic ducts to turn on a basal cell phenotype. Hence, our data suggest that published work might have missed this important cell type when using mice as an experimental model for human pancreatic physiology and disease. If basal cells were to exist in mice, using *Sox9* and *Hnf1β* as Cre-drivers may not be adequate models as human pancreatic basal cells do not express these transcription factors. *Krt19*-Cre might have been more suitable since all pancreatic basal cells, at least in humans, express KRT19.

Our findings warrant future studies on the role of pancreatic basal cells in tissue homeostasis and disease. In terms of homeostasis, a consensus exists that adult endocrine and exocrine pancreatic cells can maintain a slow turnover by low level replication. Evidence for stem cells in the adult pancreas has been highly disputed and has mostly relied on mouse modelling where *de novo* generation of insulin-producing beta-cells was examined. Only in case of substantial tissue injury, (facultative) stem cells were found to come into play (45). It is conceivable that basal cells would be a different and ‘last resort’ stem cell, reminiscent of what has been reported in the skin and the intestine, where different types of stem cells reside, confined to restricted niches within the tissue.

Here this niche seems to express markers of GI stem cells. Not all their fate-potential is called upon during physiological conditions, only specific conditions may force the cells into action (46). It may be the organoid culture conditions that provoke such response, specifically in SOX9-low expressing duct cells, suggesting that SOX9 may be a barrier to basal cell fate. Interestingly, the pancreatic ducts have often been referred to as a stem cell niche for β-cells and Dirice et al. described SOX9-negative cells located in the duct that contribute to new β-cells (47). This would fit with our findings of basal cells located in ducts with undetectable SOX9 levels.

In terms of disease, we demonstrate that the pancreatic basal cells expand significantly in chronic pancreatitis suggesting an active contribution to its pathogenesis. It remains to be investigated whether this expansion results from proliferation of pre-existing rare basal cells or recruitment of (differentiated duct) cells into a basal cell state. Early stage samples of disease that would facilitate such study are however scarce.

One clear gap of knowledge is the development of the different histologically and molecularly defined subtypes of PDAC. Ontogeny and plasticity of the subtypes could be driven by oncogenic mutations and environmental stress but could also be determined by the nature of the cells of origin. Thus far, mouse experiments have shown that pancreatic acinar and duct cells have potential to generate tumors, albeit likely upon different mutational drivers and through different precursor lesions (1,48). Evidence in patient samples is emerging that different PDAC phenotypes share epigenetic traits with the cells of origin being that the methylation and gene expression signatures of ductal cells are retained by the more aggressive basal subtype (Espinet et al., under review). With our new findings, the role of basal cells, being a subset of ductal cells, should be considered. The inverse correlation of ΔNp63 expression with SOX9 that we observe in the human samples and the mouse organoid cultures also reminds us of our previous cancer study where the squamous/basal subtype showed the lowest SOX9 expression (49). Even if the identity of the cell of origin is not a (sole) determinant of the eventual tumor subtype, a better understanding of the basal cells may provide critical insights in the development of a basal-like pancreatic tumor. Chan-Seng-Yue et al. (50) and Miyabayashi et al. (51) pointed to the possible evolution of some but not all tumors with a classical phenotype into a basal phenotype, driven by amplification of mutant *KRAS*. One could speculate that tumors arising from basal cells with stem cell features would present a wider differentiation potential, thus being more susceptible for transitioning from classical to basal than those arising from a cell in a committed differentiation state.

In conclusion, we discovered basal cells in the human adult pancreas and developed FLIP-IT, a protocol to visualize and analyze (these) cells in 3D using high resolution microscopy. We showed expansion of basal cells specifically in disease contexts. In the light of this new discovery, the established role of basal cells in other tissues and their absence in commonly used mouse experimental models, our observations compel re-interpretation of the cellular pathogenesis of pancreatic diseases.

## Acknowledgements

We would like to thank the Centre for Inflammation Research at VIB-UGent for pancreata of their transgenic mouse model, the Cell Differentiation lab for sections of neonatal mice and the use of their antibodies, the Beta Cell Neogenesis lab for sections of the PDL mice and the Reproduction and Genetics lab for the use of their antibodies. We thank the Central Biobank UZ Brussel for the human samples.

## Contributors

This study was conceptualized and designed by SM, MVB, MR and IR. SM, MVB, KC, HM, GL and MR performed experiments, data collection, and interpretation. FE, HH, LB, PI, PR and FXR provided intellectual input and important samples. SM, MVB and IR wrote the manuscript, and all authors edited the manuscript.

## Patient consent for publication

Retrospective use of patient samples was approved by the Medical Ethical commission of the UZ Brussels and the Central Biobank UZ Brussel.

## Ethics approval

Mouse experiments in the Cell Differentiation lab received ethical approved (16-277-1 (LA1230277). Ethical approval for the mouse experiments at de Duve institute received ID 2019/UCL/MD/005. Mouse experiments in the University of Pittsburgh Medical Center received ethical approval under ID 18022411.

## MATERIALS AND METHODS

### Human samples

Human samples (FFPE embedded tissue blocks and their donor characteristics) were collected from the Beta-cell bank UZ Brussels. Pancreata of Whipple resections, chronic pancreatitis and autopsy samples plus patient characteristics were obtained from the department of Anatomopathology of UZ Brussels. Ethical consent was given by the Committee of Medical Ethics - UZ Brussels and samples were obtained through the Central Biobank UZ Brussel (17-183).

### Mouse samples

Mice were sacrificed in accordance with institutional ethical guidelines and regulations and were approved by VUB Animal Ethics Committee (ethical approval 19-595-3). Mouse experiments in the Cell Differentiation lab received ethical approved (16-277-1 (LA1230277)). Ethical approval for the mouse experiments at de Duve institute received ID 2019/UCL/MD/005. Mouse experiments in the University of Pittsburgh Medical Center received ethical approval under ID 18022411.

#### Hematoxylin-eosin Saffron staining

After baking, deparaffination and rehydratation, sections were stained with hematoxylin for 5 minutes and were then differentiated with alcoholic acid and were blued in lithium carbonate. The slides were then stained with Erythrosin B for 2 minutes and 30 seconds. Slides were dehydrated in propanol and were then stained with saffron for 30 seconds and mounted with Pertex mounting medium.

### Immunohistochemistry

For singleplex ΔNp63 stainings on human samples, the automated stainer Ventana Benchmark ULTRA was used with the anti-p40 antibody, clone BC28 (790-4950, Roche, Switzerland). For brightfield doubleplex stainings, the Benchmark ULTRA was used with the anti-p40 antibody in combination with the anti-CA19.9 antibody (760-2609, Roche) and the anti-MUC6 antibody (760-4390, Roche). For these double stainings and other double stainings mentioned later, we limited our investigation to samples of chronic pancreatitis, as they contained a larger population of ΔNp63^+^ cells to characterize.

On mouse pancreas, we performed manual DAB stainings as the automated stainer detects all mouse IgGs. One to three random sections were assessed per block (S table 2). Sections were baked, deparaffinated and rehydrated. Endogenous peroxidase activity was blocked using 3% Hydrogen Peroxidase in methanol for 30 minutes. Antigen retrieval was performed using citrate buffer in a pressure cooker for 40 minutes, and protein block was done using casein block concentrated 25%. The primary antibody was incubated overnight at 4°C. The antibodies anti-p40 (ab167612, Abcam; diluted 1/200), anti-p40 (ABS552, Sigma-Aldrich; diluted 1/50.000), anti-KRT14 (1/2000, HPA023040, Atlas antibodies) or anti-KRT5 (1/200, ab52635, Abcam) were used, which gave identical results. The next day, slides were incubated for 30 minutes with biotinylated goat anti-rabbit IgG antibody (Vector, BA-1000, 1/200). Afterwards, slides were incubated for 30 minutes with a Streptavidin-biotin-HRP complex (VECTASTAIN(R) ELITE(R) ABC HRP Detection Kit, Vector). DAB incubation was done by diluting DAB concentrate 10 times in peroxide buffer (11718096001, Roche), after which slides were counterstained with haematoxylin for 30 seconds, differentiated in alcoholic acid and blued in lithium carbonate. Slides were dehydrated and mounted with Pertex mounting medium. For these stainings, a positive control of either a mouse mammary gland or mouse skin were included.

For immunofluorescence (IF), we resorted to a P63 antibody (ab735, Abcam) as the ΔNp63 antibodies mentioned before, and additionally anti-p40 (ab172731, Abcam) were not suitable for multiplex IF. Slides were processed as above and incubated overnight at 4°C with a cocktail of antibodies, always including the P63 antibody (1/50). Other primary antibodies used were E-cadherin (1/100, AB5733, Sigma-Aldrich), KRT19 (1/100, TROMA-III, DSHB), SOX9 (1/1000, ABE2868, Sigma-Aldrich), HNF1β (1/100, sc-7411, Santa Cruz), PDX1 (1/100, AF2419, RandD), KRT5 (1/100, ab52635, Abcam), KRT14 (1/1000, HPA023040, Atlas antibodies), S100A2 (1/200, HPA062451, Atlas antibodies), KRT7 (1/2000, ab181598, Abcam), DCLK1 (1/500, ab109029, Abcam), CD142 (1/100, AF2339, RandD), OLFM4 (1/5000, HPA077718, Atlas antibodies), MUC6 (1/4000, ab223846, Abcam), KI67 (1/1000, 14-5698-82, Ebioscience), NANOG (1/200, 49035, cell signaling technology) and OCT3/4 (1/200, 561556, BD Biosciences). The next day, slides were incubated with a cocktail of secondary antibodies for 1 hour. Antibodies used were donkey anti-mouse Cy3 (1/500), donkey anti-mouse AF647 (1/500), donkey anti-rabbit AF647 (1/500), donkey anti-chicken Cy3 (1/100), donkey anti-rat AF647 (1/500), donkey anti-goat AF647 (1/500) and donkey anti-rabbit AF488 (1/500), all from Jackson ImmunoResearch. After rinsing, slides were mounted with fluorescent mounting medium with DAPI added at 10 μg/ml. A section of human skin was included as positive control for ΔNp63.

#### RNA in situ hybridization (BaseScope)

For the BaseScope analysis, the standard protocol for BaseScope on FFPE tissue from ACDBio was used with the Basescope RED v2 kit. In short, 5μm slides were baked and deparaffinized, slides were incubated with hydrogen peroxide and next target retrieval was performed for 15 minutes. Protease III was applied for 15 minutes, and then the probe was incubated for 2 hours. A custom probe was designed to bind specifically to the promotor region of the delta isoform of P63, therefore not detecting the TA isoform. Standard hybridization followed, completed by signal detection and subsequent hematoxylin counterstaining. Slides were mounted with Pertex mounting medium.

### FLIP-IT

#### FFPE processing

Samples were verified for P63 presence in 2D sections. 5mm punches (up to 37 mm^3^ of tissue) were acquired with the Tissue-Tek Quick-Ray Tissue Microarray System (Sakura, Torrance, USA). The paraffin was visually eliminated from the punches using a heater and then incubated in Hemo-De (Laborimpex 23412-01) overnight at room temperature. Afterwards samples were incubated in ethanol and rehydrated. Then washed for 3 incubations in PBS-Triton 0,5%.

#### Delipidation

Samples were incubated with clearing solution for 3 days. The clearing solution was refreshed every day. The next incubation steps were performed on an orbital shaker. Samples were washed with PBS-Triton 0,5% to rinse out remaining micelles.

#### Blocking and immunostaining

Samples were blocked overnight in blocking solution with 25% casein block (Thermo, 37528).

#### Immunostaining

Samples were incubated with primary antibody for 3 days. Primary antibodies used: KRT5 (1/200, ab52635, Abcam), KRT7 (1/100, ab181598, Abcam), TROMAIII (1/50, DSHB), P63 (1/50, ab735, Abcam) and Laminin (1/25, L9393, Merck). After primary antibody incubation, samples were washed three times with washing buffer for 1 hour each and then incubated with Alexa Fluor donkey anti-mouse 647-, anti-rat Cy3-, and anti-rabbit 488-conjugated secondary antibody for 3 days. Next, samples were washed 3 times with washing buffer for 1 hour. All steps were performed at room temperature while samples were gently shaken in amber 5mL tubes (Eppendorf) to protect the samples from light.

#### Refractive index matching and agarose gel embedding of the cleared tissue

FFPE samples were incubated in 50% and 100% CUBIC-R for at least 6h each. Fresh samples did not require additional CUBIC-R incubation. 2% agarose was made by dissolving low melting point agarose powder (Sigma, A4018) in Fresh CUBIC-R in the microwave. To form the bottom gel layer (2mm height), 0.304mL of the solution was transferred with a P1000 pipette (Eppendorf) in a custom-made glass chamber, covered with a cover glass (Leica) and incubated at 4°C for 15 minutes. To form the middle gel layer, 1.5mL of the mixture was poured into a 5mL tube to which the sample was added and carefully poured in the chamber and incubated at 4°C for 30 minutes. To form the top gel layer, the remaining mixture was poured in the chamber until the surface protruded slightly, covered with a cover glass, and incubated at 4°C for at least 30 minutes. The sample was then removed from the chamber and immersed in CUBIC-R. Next, the sample was dehydrated in ascending ethanol solutions and subsequently incubated in ascending ECi solutions. The next day the sample was glued to the mount with superglue. All steps were performed while being protected from light as much as possible. The sample was immersed in fresh ECi for LSFM image acquisition.

#### LSFM Imaging

Images were acquired using a Zeiss Lighsheet Z.1 fitted with 405,488,561,638nm lasers. Samples were optically sectioned using a z-plane optimally adjusted. Overview images were acquired using 20x objective, NA=1 with zoom 0.36 – 8bit. High magnification images were acquired using 20x objective with zoom between 1 and 2,5 (magnification 20x and 50x)-16bit.

#### Dissociation and FACS analysis of pancreatic cells

Adult mouse pancreas from Sox9: eGFP transgenic mice (33) were harvested and digested in 1.4 mg/mL collagenase-P (Roche) at 37 °C for 20-30 min. Peripheral acinar-ductal units, depleted of endocrine islets, were prepared as described in (52). Following multiple washes with HBSS supplemented with 5% FBS, collagenase-digested pancreatic tissue was filtered through 600 μm and 100 μm poly-propylene mesh (BD). Peripheral acinar-ductal units containing intercalated ducts and centroacinar cells (hereinafter called small ducts of less than 100 μm), intercalated (inter- and intra-lobular ducts, hereinafter called medium ducts of 100-500 μm) and main ducts and its ramifications (hereinafter called big ducts of more than 500 μm) were further dissociated for FACS analysis in TrypLE (Invitrogen) and incubated at 37 °C for 5 min. Dispersed cells were filtered through 40μm poly-propylene mesh (BD). Dissociated cells were then resuspended at 1:106 cells per ml in HBSS supplemented with 0.5% FBS. Cell sorting was performed using a FACS-Aria II (Becton Dickinson). The sorting gate for Sox9:eGFP positive ductal cells was established by using a WT mouse pancreas sample as negative control. Cells were directly sorted in RNeasy lysis buffer (Qiagen) for RNA extraction or sorted in complete organoid media for culture.

### RNA analysis

Total RNA was isolated using the RNeasy Minikit (Qiagen) or TRizol followed by DNAse I treatment (Invitrogen) RNA was reverse transcribed with SuperScript III Reverse Transcriptase and random hexamers according to the manufacturer’s instructions. qPCR of reverse-transcribed RNA samples was performed on a 7900 Real-Time PCR system (Applied Biosystems) using the Power SYBR Green reagent (Applied Biosystems).

### Organoid culture

Entire ducts were embedded in GFR Matrigel, and cultured in organoid expansion medium (53) (AdDMEM/F12 medium supplemented with HEPES (1x, Invitrogen), Glutamax (1x, Invitrogen), penicillin/streptomycin (1x, Invitrogen), B27 (1x, Invitrogen), N-acetyl-L-cysteine (1 mM, Sigma, RSPO1-conditioned medium (10% v/v), Noggin-conditioned medium (10% v/v), epidermal growth factor (EGF, 50 ng/ml, Peprotech), Gastrin (10 nM, Sigma) and fibroblast growth factor 10 (FGF10, 100 ng/ml, Preprotech). After 3 passages we used the organoids for RNA or immunolocalization analysis.

#### Whole mount organoid staining

For whole mount organoid staining the protocol described in Dekkers et al. (54) was followed with minor modifications.

### Image processing

DAB slides were visualized and scanned with the Aperio CS2 and with the 3DHistech Pannoramic SCAN slide scanner. Slides were viewed with the Pathomation PMA.view software.

Fluorescent multiplex stainings were visualized with EVOS FL Auto Cell Imaging System. Confocal imaging was done using the ZEISS LSM 800 system. A merged Z-stack was created and saved as a PNG using the ZEISS Zen Lite program. BaseScope slides were also imaged with the ZEISS LSM 800 system, using the Cy3 channel to detect the BaseScope signal, and the brightfield channel to detect the hematoxylin staining.

Acquisitioned 3D data was processed using Zen black software using online dualside fusion algorithm. If necessary DualsideFusion files underwent background subtraction or were deconvolved using deconvolution module set to medium strength constrained iterative deconvolution. Afterwards tiled images were imported in Arivis 3.0 for stitching. 3D renderings, movies and images were acquired using Arivis software.

### Data analysis

The HALO image analysis platform was used for all quantifications of the 2D slides. Area of the tissue on each slide was quantified using the Area Quantification v2.1.3 tool. To quantify the total ΔNp63^+^ cells over all cells located in ducts, ducts were first annotated on one annotation layer and were then calculated using the ISH-IHC v1.3 quantification algorithm. To calculate the optical density of hematoxylin and eosin in ΔNp63^+^ cells and duct cells, two annotation layers were created, and a selection of ΔNp63^+^ cells and duct cells were annotated in a separate layer each. The optical density was quantified using the Area Quantification v2.1.3 algorithm. Finally, ΔNp63 expression in tumors was analyzed on tissue microarrays (TMAs), which were first segmented using the TMA module. ΔNp63 signal was quantified using the Area Quantification v1.0 algorithm.

Arivis v3.2 was used to analyze 3D high resolution images. Voxel operations, membrane segmentation and 3D object building pipelines were created in Arivis for identification of KRT7^+^ and KRT5^+^ cells while excluding cells touching the edges. Membrane segmenter was set to plane-wise segmentation allowing holes and full connectivity in X/Y/Z. The segments were interrogated for Sphericity (3D roundness/shape) and volume. Manual quality control was performed for omittance of false-positive and -negative results.

### Statistical analysis

Experimental data were analyzed by two-tailed unpaired Student t test, unpaired t test with Welch’s correction, paired t-test, Mann–Whitney or one-way Anova with Turkey’s multiple comparisons test using GraphPad Prism8.0 and statistical significance was accepted at P < 0.05. The results are shown as mean ± standard error of mean (SEM). The number of independent experiments (n) is indicated in the figure legends.

**S Figure 1: ΔNp63 expression in pancreatic ductal adenocarcinoma samples.** (A) Quantification of ΔNp63 expression displayed as optical density for four different groups of tumors (n=141), expressing no ΔNp63 (negative), only in a few cells (partially positive) and samples that express ΔNp63 (positive). Four adenosquamous sample, which all fall in the positive group, are indicated in orange. Mean ± SEM is shown. (B) Visualization of the quantification through HALO. DAB stainings are quantified in red (haematoxylin) and green (ΔNp63). (C-F) Representative images of the four different groups. (C) Negative tumor; (D) Partially positive tumor, showing basal cells in one duct; (E) Positive tumor and (F) Adenosquamous tumor.

**S Figure 2: RNA detection of ΔNp63 using BaseScope RNA *in situ* hybridization.** (A-C) RNA detection in healthy donor pancreas (A), chronic pancreatitis (B) and normal adjacent to PDAC area (C), with the corresponding P63 staining below. RNA is visualized as red dots. (D) RNA detection in a positive control tissue, human skin. (E) Validation of P63 antibody in immunofluorescence (IF) on the right with ΔNp63 antibody staining in immunohistochemistry (IHC) on the left.

**S Figure 3: ΔNp63 expression in normal pancreas is not associated with socio-demographic features of donors.** Characteristics of all human pancreas donors with ΔNp63 detected in a section (n=53) and without ΔNp63 detected in a section (n=61). (A) Age, (B) Gender, (C) Days spent in the intensive care unit and (D) BMI. Mean ± SEM is shown.

**S Figure 4: ΔNp63^+^ cells are epithelial cells.** IF for E-cadherin (green) and P63 (red).

**S Figure 5: ΔNp63^+^ cells are not located in pancreatic ductal glands.** (A) IF for P63 (red) and MUC6 (white). White arrow indicates ΔNp63^+^ cell, the orange arrow indicates a MUC6^+^ cell. (B) IHC for ΔNp63 (brown) and MUC6 (red). (C) IF for 63 (red) and KI67 (white). White arrow indicates ΔNp63^+^ cell, the orange arrow indicates a KI67^+^ cell.

**S Figure 6: ΔNp63^+^ cells have a paler cytoplasm and nucleus compared to ductal cells.** (A) HES staining of a duct in a human healthy pancreas. Inset on the right bottom shows magnification of one ΔNp63^+^ cell. Black arrows point to ΔNp63^+^ cells. (B) Consecutive slide of A showing the DAB staining for ΔNp63. Quantification of the (C) hematoxylin and (D) eosin positivity in ΔNp63 cells compared to ductal cells (n=8). One line indicates one slide that was analyzed for both ΔNp63^+^ and ΔNp63^−^ cells. (***p < 0.001, **** p < 0.0001)

**S Figure 7: ΔNp63+ cells are not myoepithelial cells.** (A) IHC for ΔNp63 (brown) and calponin (red) in a duct positive for ΔNp63. (B) shows positive control for calponin in the wall of a blood vessel.

**S Figure 8: ΔNp63^+^ cells do not express typical pluripotent stem cell markers.** (A) IF for P63 (red) and NANOG (white). Positive expression of NANOG is shown in a seminoma in panel (B). (C) IF for P63 (red) and OCT4 (white). Positive expression of OCT4 is shown in a seminoma in panel (D).

**S Figure 9: FLIP-IT overview of human and mouse sample processing.** (A) FLIP-IT protocol steps in archival FFPE human samples and representative pictures of the samples. Scale bars correspond to 2mm. (B) Table comparing key protocol steps for 3D human pancreas sample processing workflow. For FFPE samples, the complementary effect of CUBIC-R and FDA-approved optical clearing agent ECi allowed safe handling, versatile imaging, storage, transport and retention of fluorescence intensity over longer periods of time. Hong et al. were able to clear thick (0.5cm) fresh/FFPE human pancreatic samples using a modified iDISCO method. They used liquid dibenzyl ether (DBE) which we refrained from since dipping high magnifying objectives in DBE could dissolve the glues within the objectives. Moreover, the iDISCO protocol induced sample shrinkage which to some extent would distort our assessments. Hence, we devised a novel methodology, FLIP-IT, that yielded consistently transparent pancreas samples ready for LSFM within two weeks, whether applied to fresh or FFPE stored specimens (some over 25 years old) and without sample shrinkage. The duration does not take into account required imaging time which is considerably longer for confocal microscopy. (C) FLIP-IT in fresh PFA-fixed mouse samples and representative pictures of the samples. Scale bars correspond to 5mm. (D) Table comparing key protocol steps for whole mouse pancreas sample processing workflow. Messal et al. used methyl salicylate for RI-matching again dissolving glues when using dipping objectives. Dipping objectives are required for high resolution light sheet fluorescence miscroscopy. The duration does not take into account required imaging time which is considerably longer for confocal microscopy.

**S Figure 10: KRT5^+^ cell sphericity and volume changes in chronic pancreatitis compared to normal human pancreas.** (A) Sphericity quantification of KRT5^+^ and KRT7^+^ cells in both normal human pancreas and chronic pancreatitis. (B) Volume quantification of KRT5^+^ cells in both normal human pancreas and chronic pancreatitis. (****p < 0.0001, n=1, >250 cells counted per group.)

**The ΔNp63 and KRT14 antibodies showed strong positivity in positive control mouse tissues.** (A) ΔNp63 IHC staining in a healthy mouse mammary gland, staining the myo-epithelial cells. (B) ΔNp63 IHC staining in healthy mouse skin, staining nuclei in the epidermis. (C) ΔNp63 IHC staining in a human adenosquamous tumor. (D) KRT14 IHC staining in the hair follicles of a human skin section. (E) P63 IF staining of nuclei of the basal cells in the epidermis of human skin.

**S Figure 12: FLIP-IT applied to whole mouse pancreas and attached duodenum and spleen.** (A) Processing protocol of fresh mouse samples. (B) Overview 3D rendering of normal mouse pancreas stained for KRT5 (pink) and KRT7 (cyan). No KRT5^+^ were seen in the mouse pancreas. Some pink color is present in areas showing nonspecific staining (confirmed at higher magnification). Asterisk shows large duct. White dotted line shows duodenum. Yellow dotted line shows spleen. Objective 5x, zoom 0.36. Scale bar corresponds to 1mm. n=3

**S Table 1: Donor characteristics.** The table shows whether ΔNp63 expression was found, the age and sex of the donor, weight, height and BMI. Days spent in hospital before organ donation and preservation time of the pancreatic sample are also indicated.

**S Table 2: Single cell sequencing datasets investigated for expression of ΔNp63.** Species of the study Is indicated in the table. ΔNp63 expression was absent in all studies except for the study of Segerstolpe.

