## Supplemental Figures for "Discovery and 3D imaging of a novel ΔNp63-expressing basal cell type in human pancreatic ducts with implications in disease"

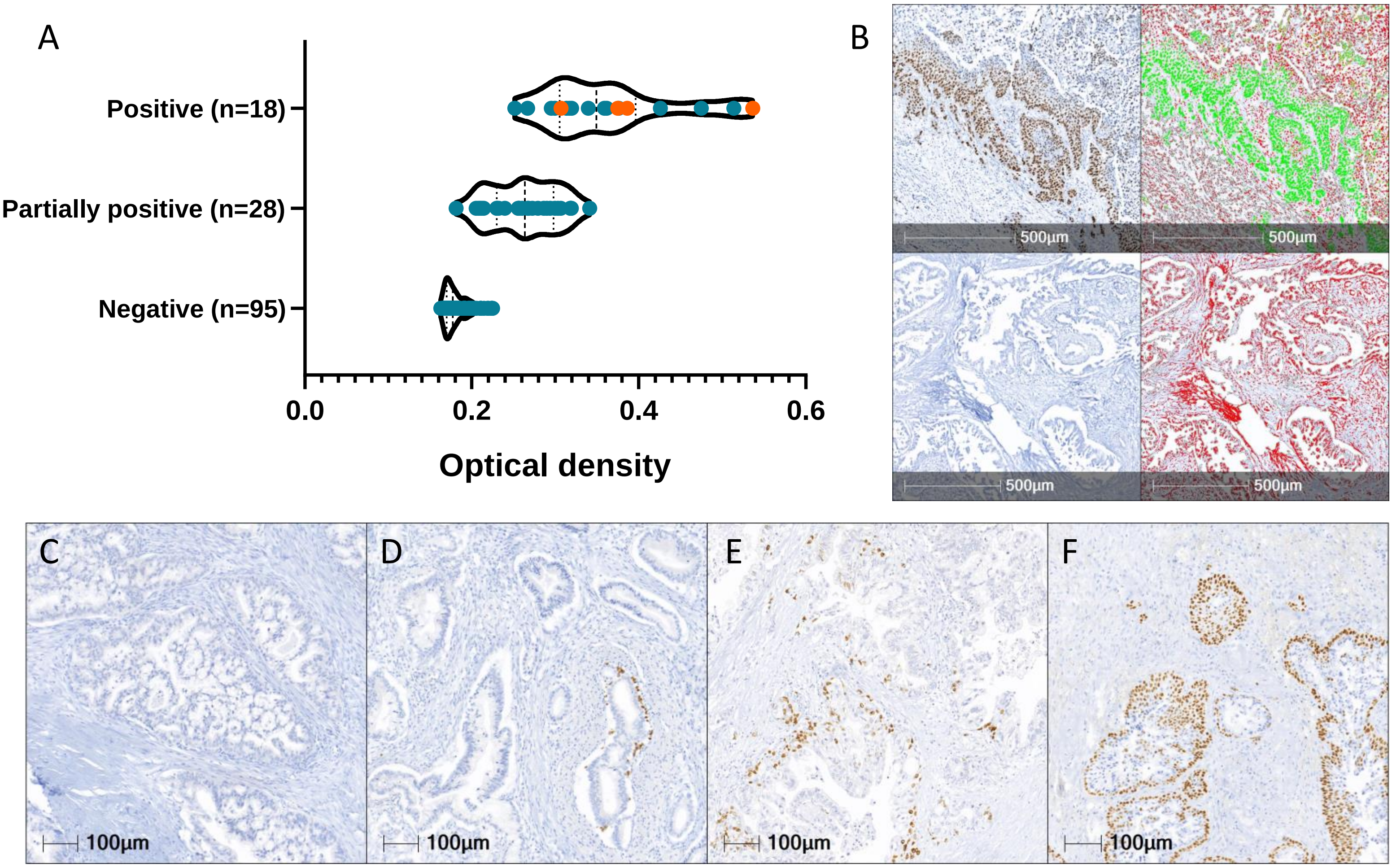
A
Positive (n=18)
Partially positive (n=28)
Negative (n=95)
0.0
0.2
0.4
0.6
Optical density
B
500µm
500µm
500µm
500µm
C
100µm
D
100µm
E
100µm
F
100µm

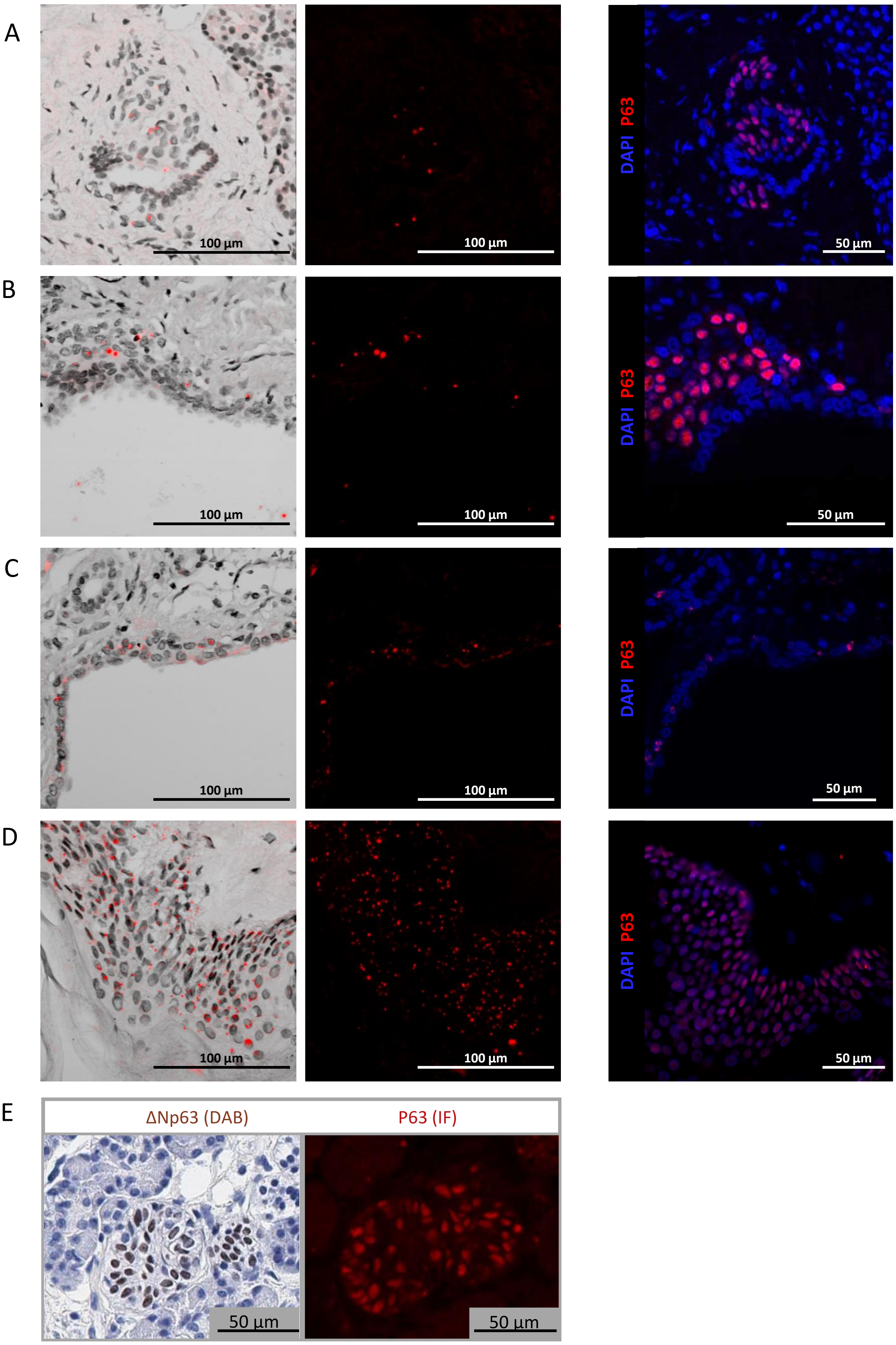
A
100 µm
100 µm
DAPI P63
50 µm
B
100 µm
100 µm
DAPI P63
50 µm
C
100 µm
100 µm
DAPI P63
50 µm
D
100 µm
100 µm
DAPI P63
50 µm
E
ΔNp63 (DAB)
P63 (IF)
50 µm
50 µm

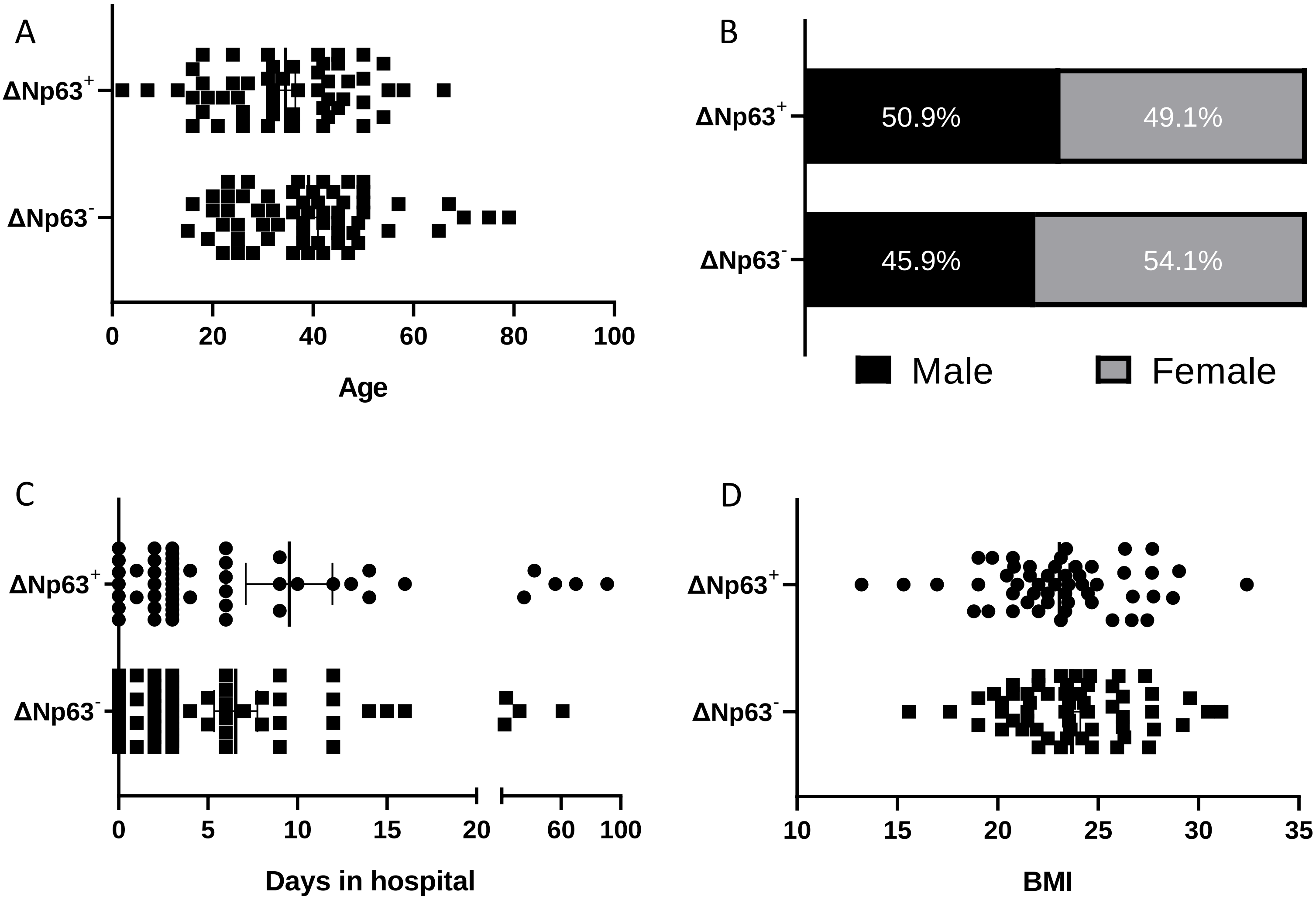

A
ΔNp63+
ΔNp63-
0 20 40 60 80 100
Age
B
ΔNp63+
ΔNp63-
50.9%
49.1%
45.9%
54.1%
Male
Female
C
ΔNp63+
ΔNp63-
0 5 10 15 20 60 100
Days in hospital
D
ΔNp63+
ΔNp63-
10 15 20 25 30 35
BMI

A

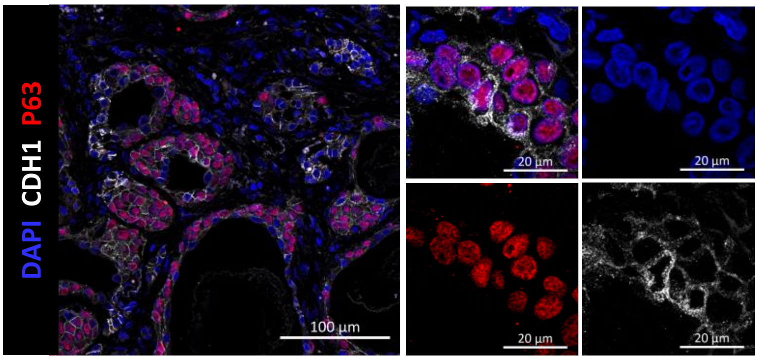

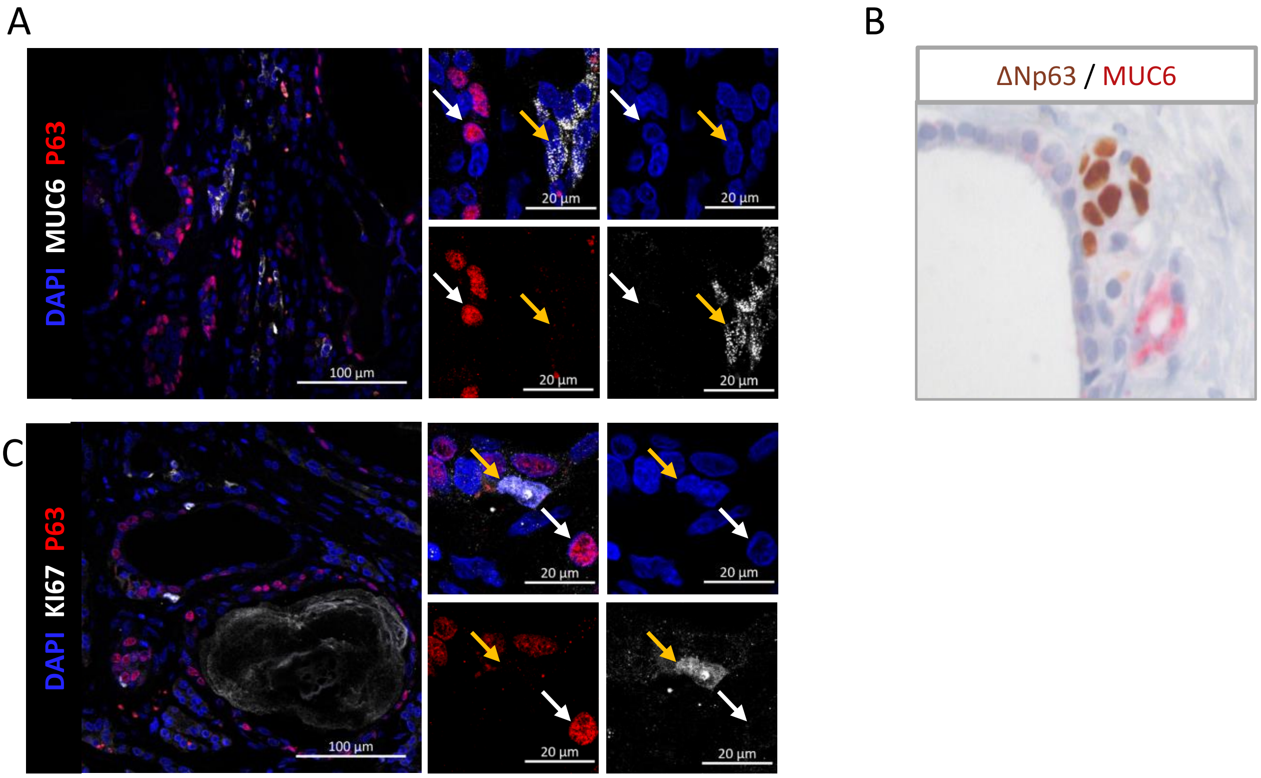
A
B
C
DAPI MUC6 P63
20 µm
100 µm
ΔNp63 / MUC6
DAPI KI67 P63

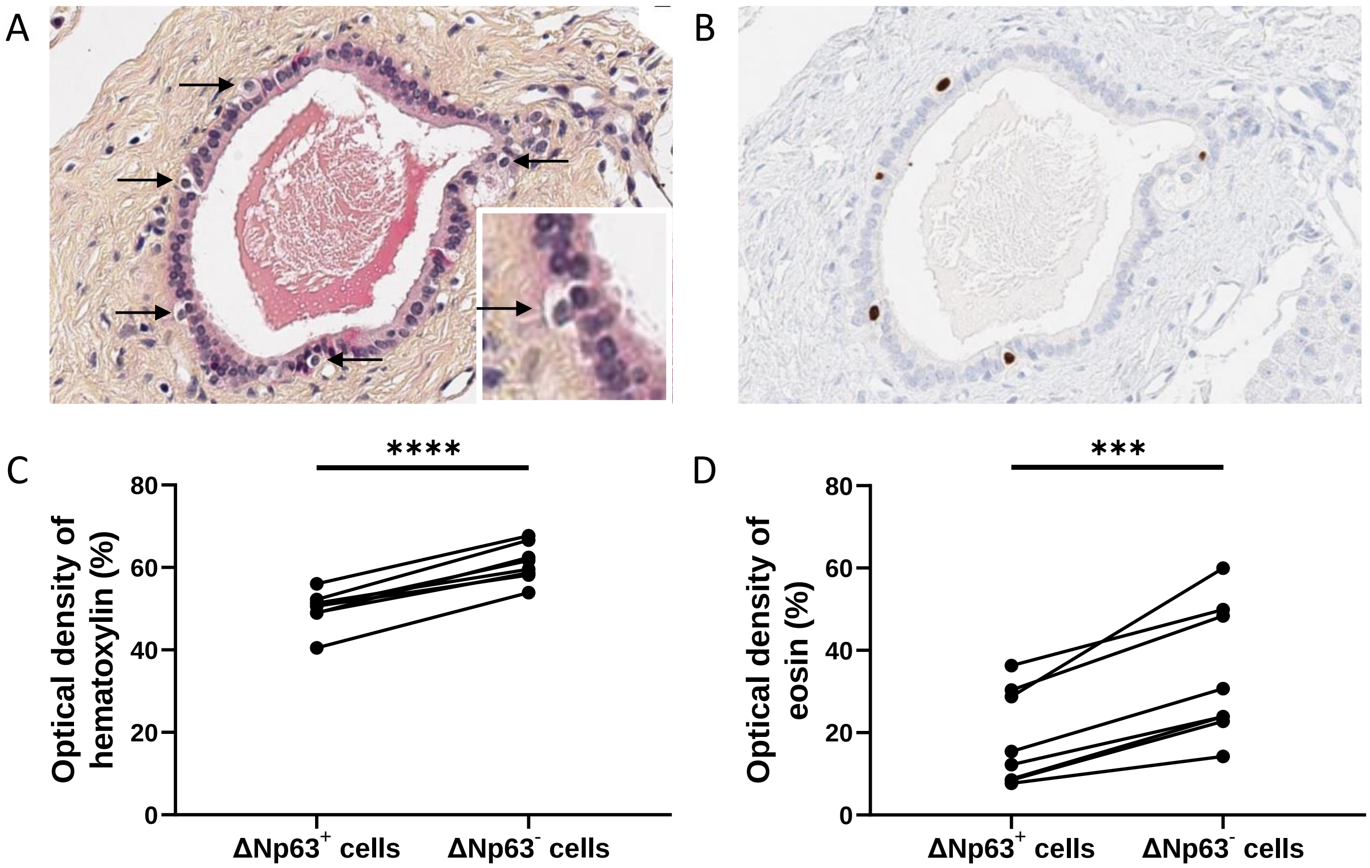
A
B
C
D
****
***
Optical density of hematoxylin (%)
Optical density of eosin (%)
0
20
40
60
80
ΔNp63+ cells
ΔNp63- cells

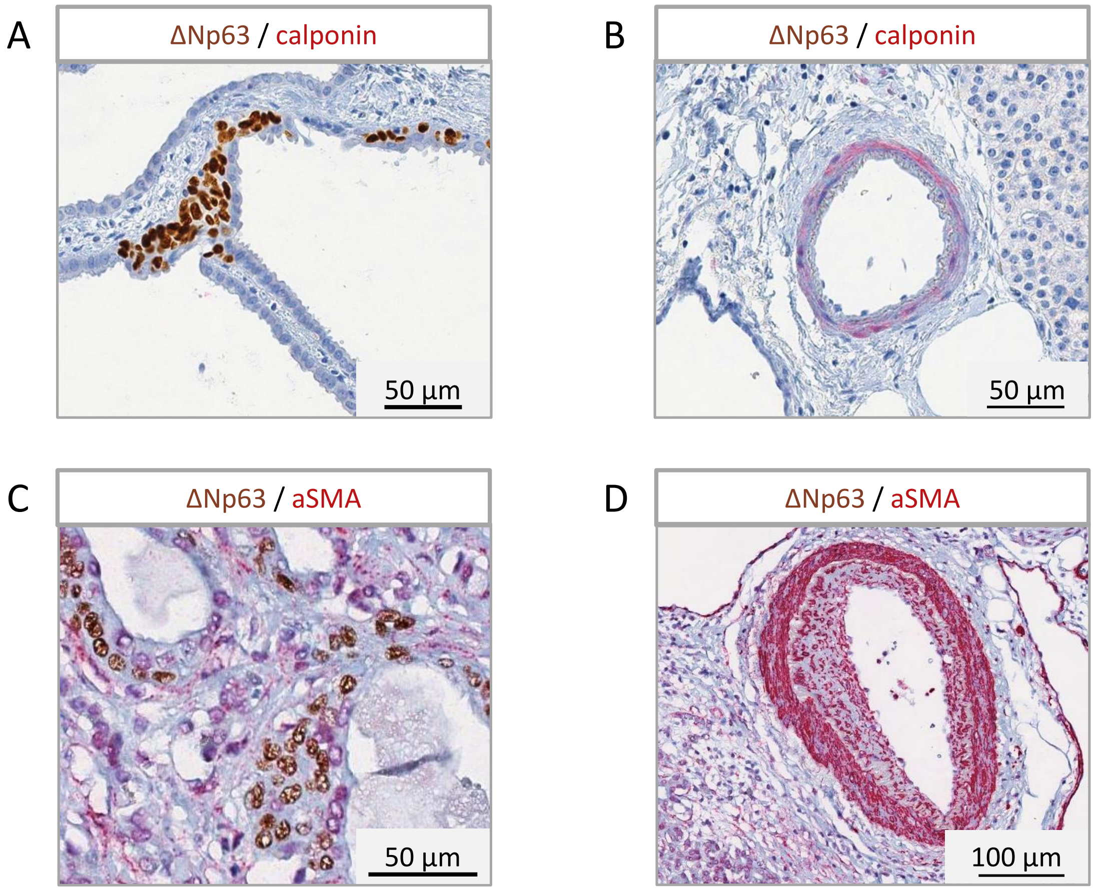

A
ΔNp63 / calponin
50 µm
B
ΔNp63 / calponin
50 µm
C
ΔNp63 / aSMA
50 µm
D
ΔNp63 / aSMA
100 µm

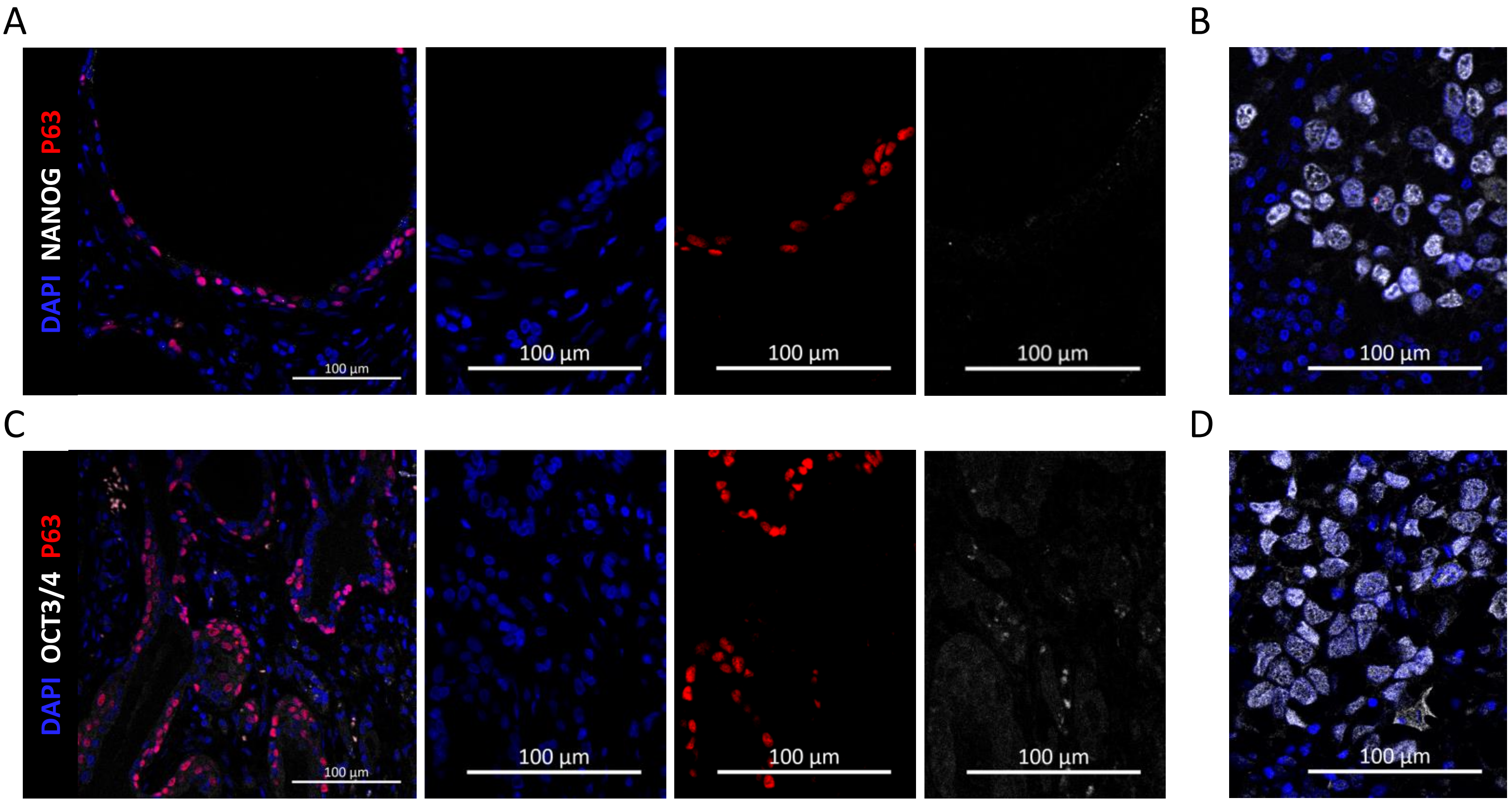
A
B
C
D
DAPI NANOG P63
DAPI OCT3/4 P63
100 µm

A

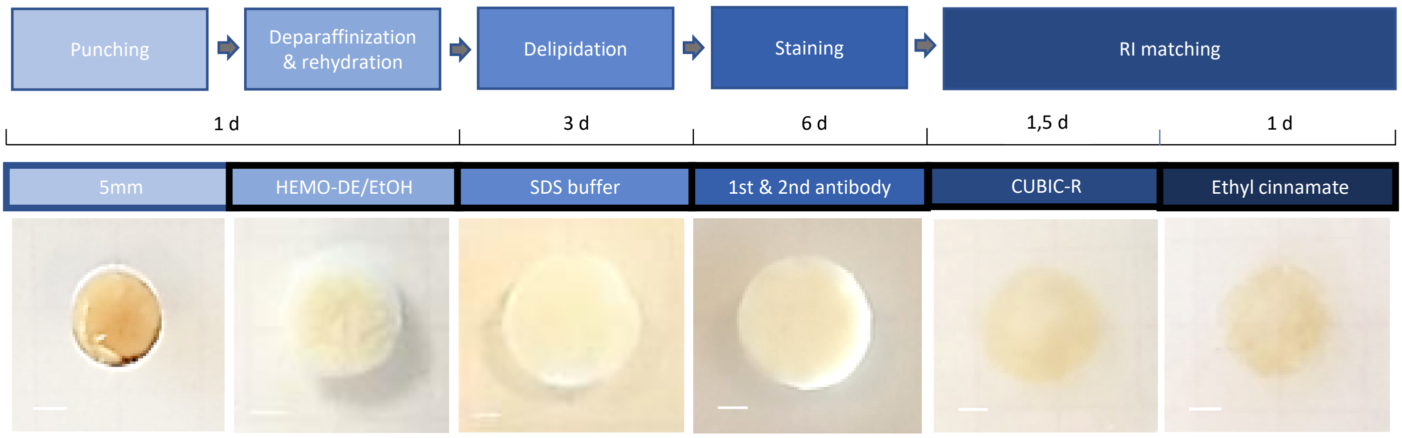


B

| Protocol | Delipidation | RI matching | Duration | High resolution imaging |
|---|---|---|---|---|
| Modified IDISCO[47] | dichloromethane/methanol | dibenzyl ether (DBE) | 20 days | confocal microscopy |
| FLIP-IT | SDS buffer | CUBIC-R/Ethyl Cinnamate(Eci) | 13 days | Lightsheet miscroscopy |

C

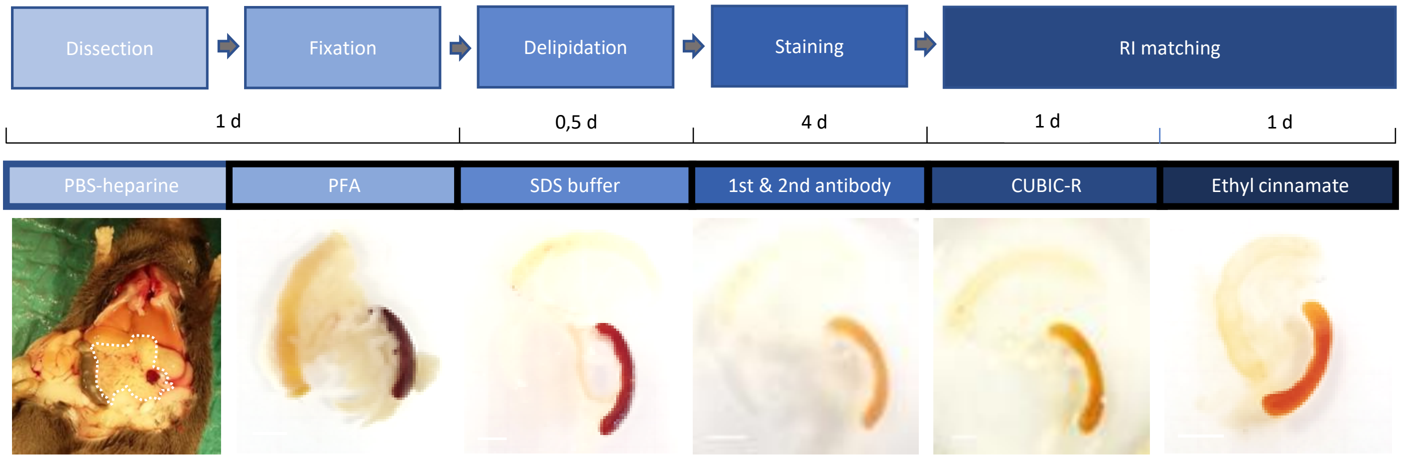


D

| Protocol | Delipidation | RI matching | Duration | High resolution imaging |
|---|---|---|---|---|
| FLASH[18] | SDS/boric acid | Methyl Salicylate | 7 days | confocal microscopy |
| FLIP-IT | SDS buffer | CUBIC-R/Ethyl Cinnamate(Eci) | 8 days | Lightsheet miscroscopy |

A

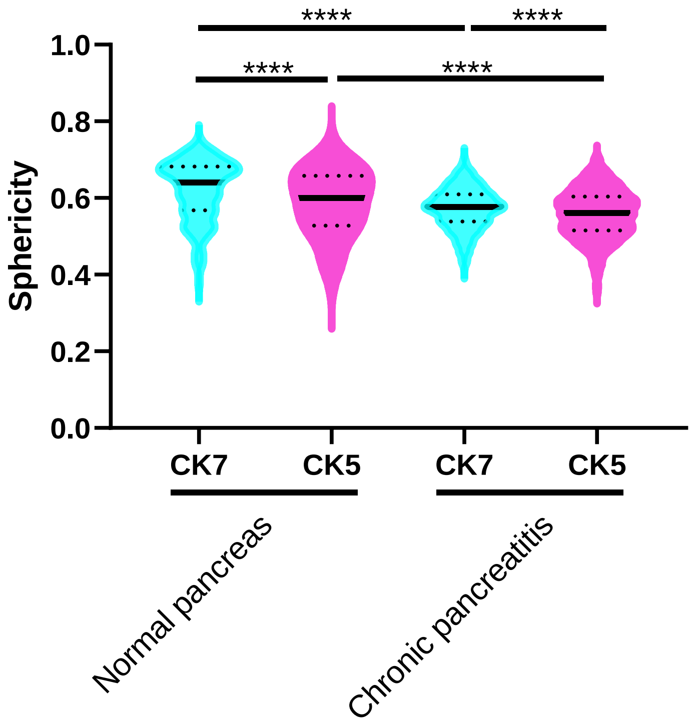

****
****
****
****
1.0
0.8
0.6
0.4
0.2
0.0
Sphericity
CK7
CK5
CK7
CK5
Normal pancreas
Chronic pancreatitis


B

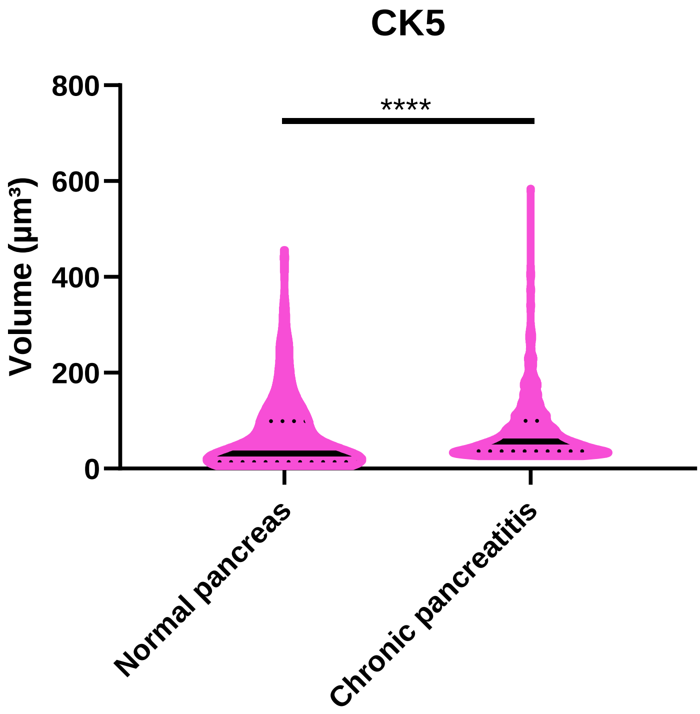

CK5
****
800
600
400
200
0
Volume (µm³)
Normal pancreas
Chronic pancreatitis

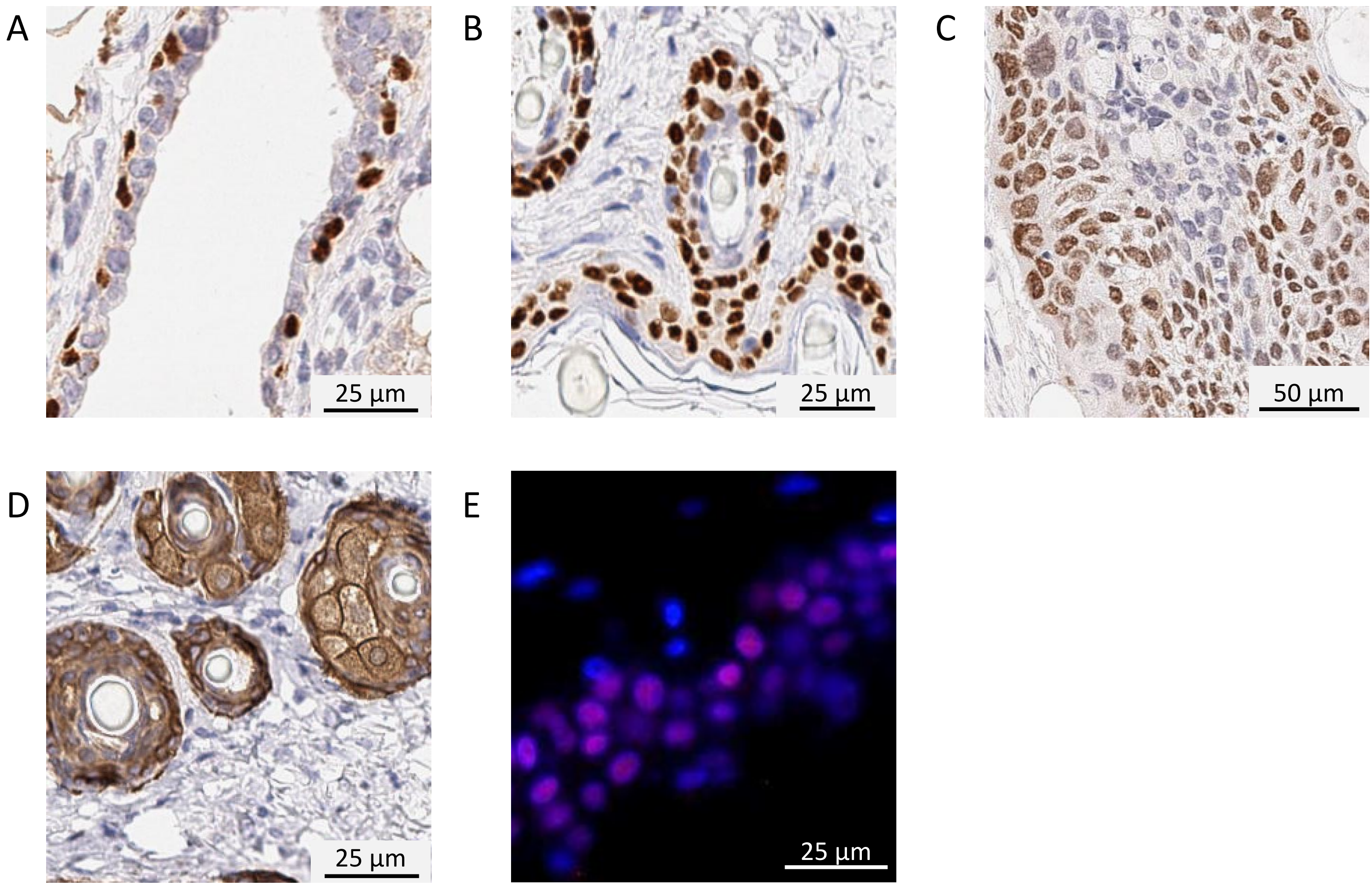
A
25 µm
B
25 µm
C
50 µm
D
25 µm
E
25 µm

A

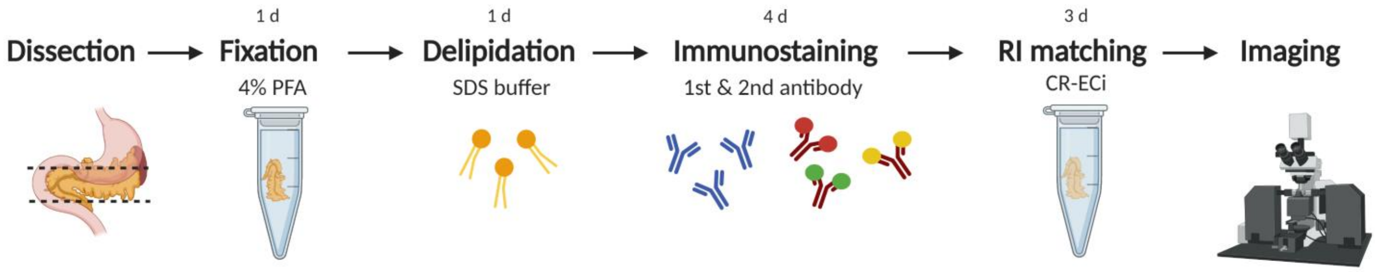
1 d
1 d
4 d
3 d
Dissection
Fixation
4% PFA
Delipidation
SDS buffer
Immunostaining
1st & 2nd antibody
RI matching
CR-ECi
Imaging

B

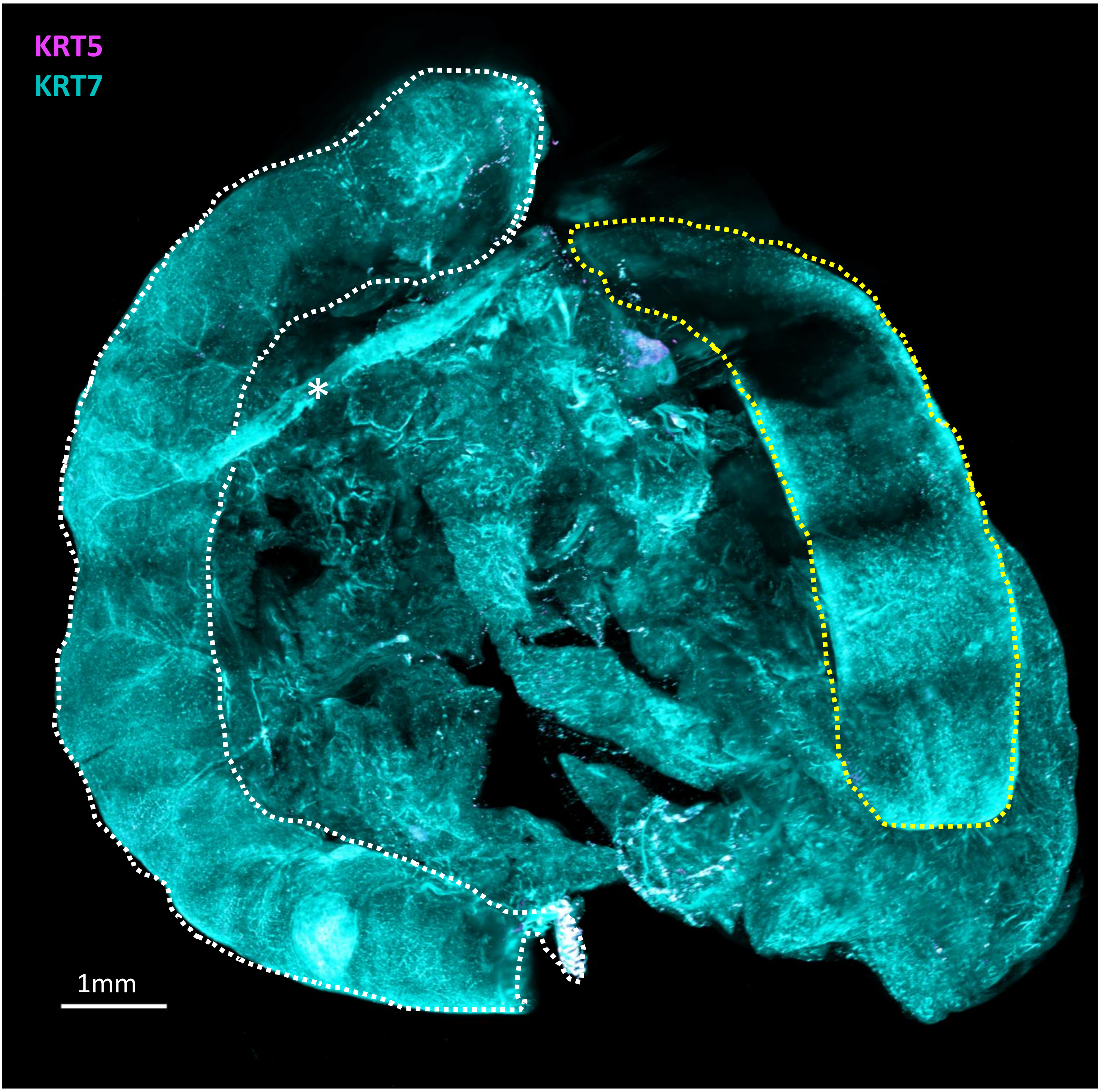
KRT5
KRT7
*
1mm
