## Supplemental Table 2 for "Discovery and 3D imaging of a novel ΔNp63-expressing basal cell type in human pancreatic ducts with implications in disease"

| **Paper** | **Species** | **ΔNp63 expression?** |
| --- | --- | --- |
| Qadir et al. (2020) | Human | No |
| Tabula Muris (2018) | Mouse | No |
| Segerstolpe et al. (2016) | Human | Yes, rarely in diabetic samples |
| Muraro et al. (2016) | Human | No |
| Enge et al. (2017) | Human | No |
| Tarifeño-Saldivia et al. (2017) | Zebrafish | No |
| Sznurkowska et al. (2018) | Mouse | No |
| Grün et al. (2016) | Human | No |
